# Molecular architecture of the luminal ring of the *Xenopus laevis* nuclear pore complex

**DOI:** 10.1101/2020.03.27.009381

**Authors:** Yanqing Zhang, Sai Li, Chao Zeng, Gaoxingyu Huang, Xuechen Zhu, Qifan Wang, Kunpeng Wang, Qiang Zhou, Wusheng Zhang, Guangwen Yang, Minhao Liu, Qinghua Tao, Jianlin Lei, Yigong Shi

## Abstract

Nuclear pore complex (NPC) mediates the flow of substances between the nucleus and cytoplasm in eukaryotic cells. Here we report the cryo-electron tomography (cryo-ET) structure of the luminal ring (LR) of the NPC from *Xenopus laevis* oocyte. The observed key structural features of the LR are independently confirmed by single-particle cryo-electron microscopy (cryo-EM) analysis. The LR comprises eight butterfly-shaped subunits, each containing two symmetric wings. Each wing consists of four elongated, tubular protomers. Within the LR subunit, the eight protomers form a Finger domain, which directly contacts the fusion between the inner and outer nuclear membranes, and a Grid domain, which serves as a rigid base for the Finger domain. Two neighbouring LR subunits interact with each other through the lateral edges of their wings to constitute a Bumper domain, which displays two major conformations and appears to cushion neighbouring NPCs. Our study reveals previously unknown features of the LR and potentially explains the elastic property of the NPC.

## INTRODUCTION

A hallmark of all eukaryotic cells is the nuclear envelope (NE), which separates the nucleoplasm, where the genetic material is stored, away from the cytoplasm, where nuclear-transcribed RNA is translated into protein of diverse functions. Nucleocytoplasmic shuttling of all substances needed for transcription and translation and numerous other cellular processes is mediated by the nuclear pore complex (NPC)^1-3^. NPC associates with and stabilizes a highly curved section of the NE – namely the fusion between the inner (INM) and outer nuclear membranes (ONM)^4,5^. NPC is among the largest supramolecular complexes in cells, with a combined mass of approximately 50 MDa in yeast^6,7^ and 110-125 MDa in higher eukaryotes^3,8-11^. The protein components of NPC are known as nucleoporin (Nup). An NPC has about 34 different Nups, most of which are conserved among different organisms, and each Nup is represented in multiple copies^11^.

X-ray structures have revealed a wealth of information on individual components and subcomplexes of the NPC^11^. This information, together with EM and other studies, have yielded a three-dimensional model of the NPC. Cryo-ET reconstruction of the NPC has been achieved at average resolutions of 58 Å, 28 Å, 30 Å, 20 Å, and 23 Å, respectively, for *Dictyostelium discoideum* (*D. discoideum*)^12^, *Saccharomyces cerevisiae* (*S. cerevisiae*)^7^, *Chlamydomonas reinhardtii*^13^, *Xenopus laevis* (*X. laevis*)^14^, and *Homo sapiens* (*H. sapiens*)^15^. NPC has an eight-fold symmetry along the nucleocytoplasmic axis and a pseudo two-fold rotational symmetry in the plane of the NE. The scaffold of an NPC is proposed to comprise a cytoplasmic ring (CR), an inner ring (IR) and a nuclear ring (NR)^5^. Cytoplasmic filaments and nuclear basket are attached to the CR and NR, respectively^5,10^.

The luminal ring (LR), as the name indicates, resides in the lumen of the NE and surrounds the NPC at the site of membrane fusion^16,17^. The LR is separated from all other components of the NPC by the nuclear membrane and may play a role in anchoring the NPC to the NE^16^. Previous EM and ET studies have revealed few describable features of the LR^12,14,16-20^ except that the LR of the yeast NPC was found to comprise eight circumferential arches^6,7^. At present, the overall organization, structural features and functional mechanism of the LR remain largely enigmatic.

Due to its location, the LR is speculated to be composed of integral membrane proteins. Among the vertebrate Nups, only four have been found to be integral membrane proteins: GP210 (Pom152 in yeast) with a single transmembrane helix (TM)^21-23^, POM121 with a single TM^24^, NDC1^25,26^ and TMEM33 (ref. 27) each with six predicted TMs. GP210 is the only Nup that contains a sufficiently large luminal domain in vertebrates for the formation of a ring scaffold in the lumen^17,28,29^.

Here we report the cryo-ET and cryo-EM structures of the NPC from *X. laevis* oocyte, which reveal elaborate structural features of the LR. These features may define and potentially explain the functions of the LR.

## RESULTS

### Overall structure of the NPC from *X. laevis* oocyte

For each *X. laevis* oocyte, the NE was isolated and transferred onto an EM grid. The grid was plunge-frozen and imaged on a Titan Krios microscope. To minimize the effect of preferred sample orientation, 1,575 tilt-series were recorded using a combination of continuous-, bidirectional- and dose-symmetric^30^ schemes, each from −60° to 60° with a 3°-increment using SerialEM^31^ (Supplementary information, Fig. S1). 36,529 NPC particles were averaged to generate a reconstruction with 8-fold symmetry using Dynamo^32^. For these particles, individual subunits of the CR, IR, NR, and LR were sub-boxed and subjected to three-dimensional (3D) classification using RELION3.0 (ref. 33) (Supplementary information, Fig. S2). Analysis by sub-tomogram averaging (STA) led to reconstruction of the CR, IR, NR, and LR at average resolutions of 9.1 Å, 13.1 Å, 13.6 Å, and 15.1 Å, respectively (Supplementary information, Fig. S3a-d). For the purpose of display in the original tomogram, 780 ribosomal particles were also subjected to the STA procedure, yielding a reconstruction at 16.4 Å resolution.

Cryo-ET allows direct visualization of all macromolecular complexes on the original tomogram of the *X. laevis* NE. In the luminal regions that surround the periphery of the NPC, additional densities are clearly visible and appear to form arch-shaped repeating structures (Fig. 1a). These densities are thought to come from the LR^16,19^. Based on the positional coordinates, reconstructions of the CR, IR, NR, LR and ribosomal subunits were individually projected back into the original tomograms (Fig. 1b and Supplementary information, Video S1). Examination of four evenly-spaced layers of the same region of a tomogram along the nucleocytoplasmic axis reveals spatial organization of the CR, IR, NR, LR, and the ribosomes (Fig. 1c and Supplementary information, Fig. S4). Nearly every NPC is surrounded by other NPCs and the ribosomes. The LR density encircles the NPC and fills the space between the INM and ONM (Fig. 1b, c).

**Fig. 1.**
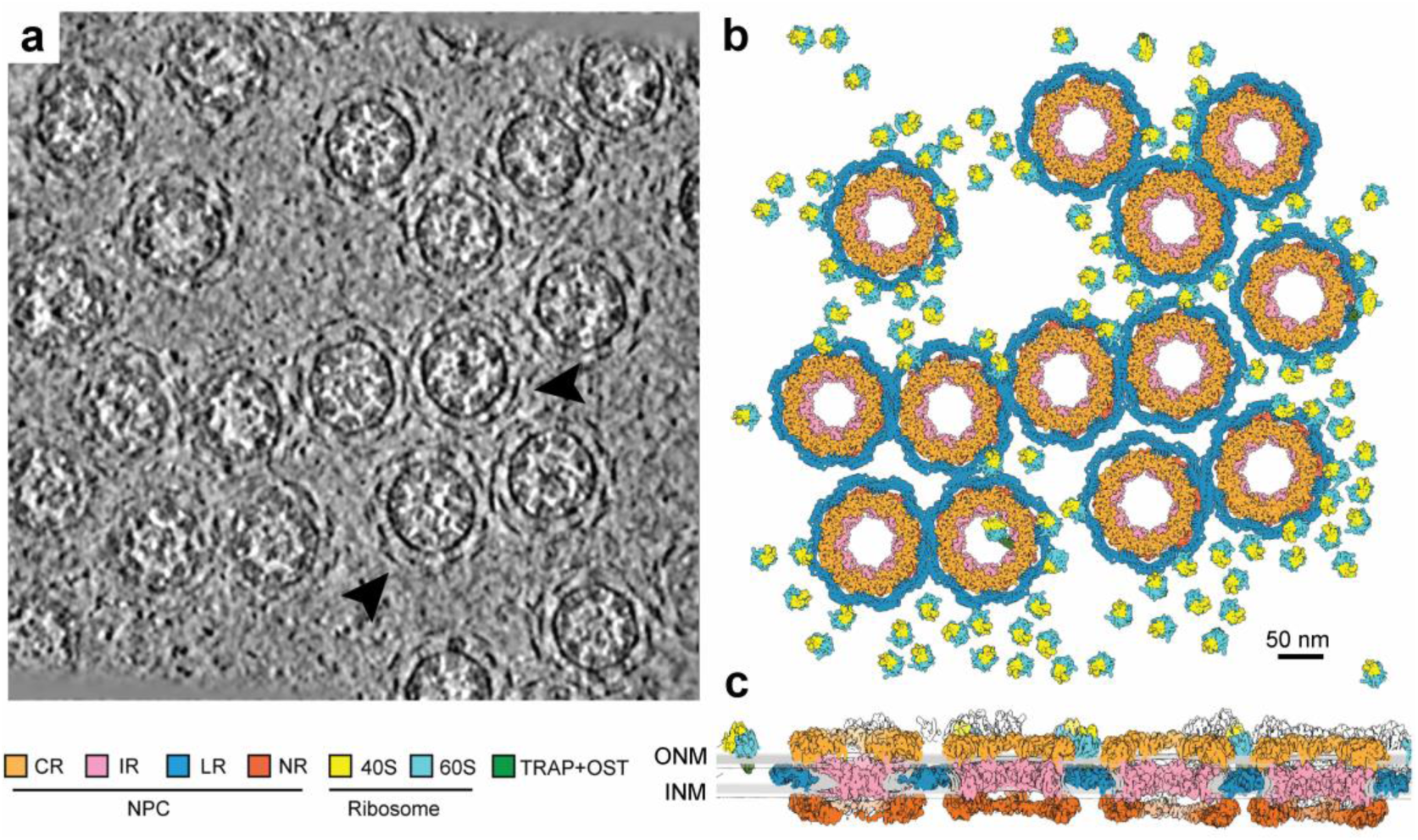
Three-dimensional organization of the NPCs in a local region of the X. *laevis* NE. **a** Organization of the NPCs in an original tomogram slice as viewed along the nucleocytoplasmic axis. Some of the representative array-like densities of the LR are indicated by arrowheads. The thickness of the tomographic slice shown here is 8.9 Å. **b** Organization of the NPCs in the reconstructed tomogram as viewed along the nucleocytoplasmic axis. As the outer boundary of the NPC, the LR appears to cushion the contacts among neighbouring NPC particles. Reconstructions for the individual NPC subunits (CR, IR, NR and LR) and the ribosomes associated with TRAP+OST^46^ were back-projected onto the original tomograms based on the refined coordinates of the individual particles. Shown here is a section of the NE from panel **a**. Scale bar, 50 nm. **c** Organization of the NPCs in the reconstructed tomogram as viewed perpendicular to the nucleocytoplasmic axis. In contrast to other ring scaffolds of the NPC, the LR resides in the lumen. 40S: Small ribosome subunit; 60S: Large ribosome subunit; TRAP: translocon-associated protein complex; OST: oligosaccharyl transferase.

Our reported average resolutions display directional anisotropy, particularly along the Z-axis (Supplementary information, Fig. S3c). Such anisotropy may reduce confidence on the interpretation of detailed structural features. To address this issue and to validate the cryo-ET STA reconstruction, we determined the cryo-EM structure of the LR using a completely independent data set through the single particle analysis (SPA) approach. During cryo-EM data collection, the sample grids were tilted at fixed angles of 0, 30, 45 and 55 degrees, generating 12,399 good micrographs^34^. The SPA approach resulted in the reconstruction of the LR subunit at an average resolution of 10.7 Å (Supplementary information, Figs S5, S6). The overall architecture and organization of the LR are nearly identical between the STA and SPA reconstructions (Fig. 2a,b). In particular, the key structural features of the STA reconstruction of the LR subunit can be very well super-imposed to those of the SPA reconstruction (Fig. 2c).

**Fig. 2.**
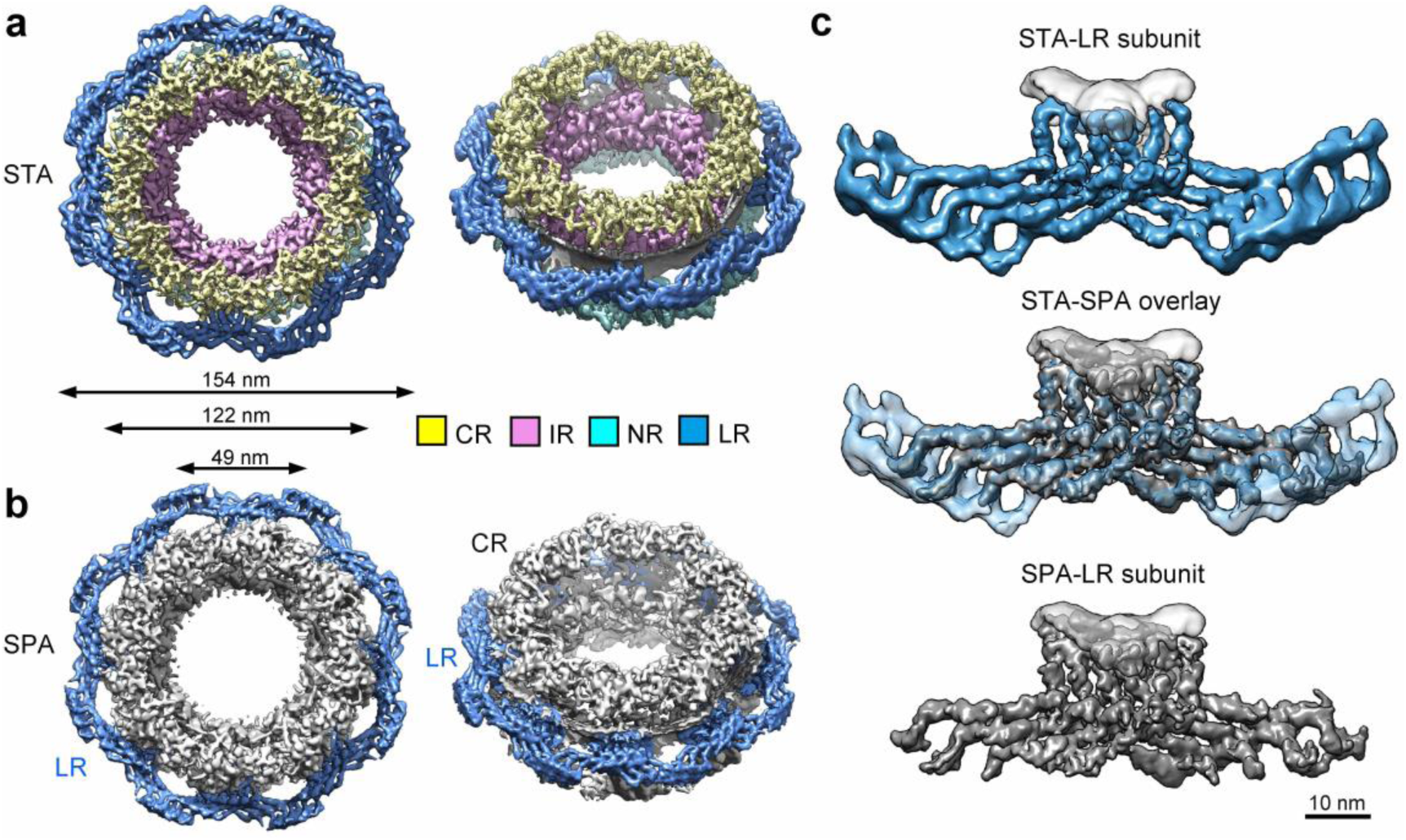
Key features of the LR in the cryo-ET reconstruction are confirmed by an independent cryo-EM reconstruction of the *X. laevis* NPC. **a** Reconstruction of a representative NPC particle by sub-tomogram averaging (STA). A top view and a tilt-45° view are shown. The individual subunits of the CR, IR, NR, and LR were projected back into the original tomograms to allow reconstruction of a number of NPC particles. Shown here is a representative NPC particle, which contains four ring scaffolds: CR (colored yellow), IR (pink), NR (cyan), and LR (marine). Viewed perpendicular to the NE (left panel), the IR and LR define the inner and outer diameters, respectively, of the cylindrical NPC. **b** Reconstruction of the NPC by single particle analysis (SPA). CR, IR and NR were reconstructed using the C8 symmetry. The LR was reconstituted using the refined LR subunit and the C8 symmetry. The LR is highlighted. **c** The key structural features of the LR subunit are nearly identical between the STA reconstruction (marine, top panel, accession code EMD-0983) and the SPA reconstruction (grey, bottom panel, accession code EMD-0982). Their overlay is shown in the middle panel. Scale bar, 10 nm.

One representative *X. laevis* NPC measures 49 nm in inner diameter (Fig. 2a,b). The outer diameter is approximately 122 nm without the LR and 154 nm with the LR. The outer boundary of the cylindrical NPC is defined by the LR (Fig. 2a,b), which is separated from the other three ring scaffolds by the nuclear membrane. The overall size of the NPC described in our study resembles that reported for two vertebrate NPCs, one also from *X. laevis*^14^ and the other from *H. sapiens*^15^, but contrasts that of the *S. cerevisiae* NPC^7^ (Supplementary information, Fig. S7). The cylindrical height of the reconstructed NPC shows some variations among the four representative organisms. Due to differences in resolution and sample preparation, detailed structural comparison of the NPC in different organisms should be performed with caution.

### Structure of the LR

Structural features of the LR subunit are reported in the main text for the SPA reconstruction (Fig. 3). Nearly identical features are shown in the supplemental information for the STA reconstruction (Supplementary information, Fig. S8). The LR has eight subunits, each comprising two symmetric wings (Fig. 3a,b and Supplementary information, Fig. S8a,b). The two wings span a distance of 70 nm and interact with each other through an extended interface (Fig. 3b and Supplementary information, Fig. S8b). Each wing contains four parallel, planar-arranged, elongated protomers (Fig. 3c and Supplementary information, Fig. S8c). Each protomer consists of an arm at one end, a central hub, and a leg at the other end. Within the same LR subunit, two wings interact with each other mainly through their eight hubs, generating a rigid structure that is termed the Grid domain (Fig. 3b and Supplementary information, Fig. S8b). These two wings also form an interface through their eight arms, which together constitute the Finger domain.

**Fig. 3.**
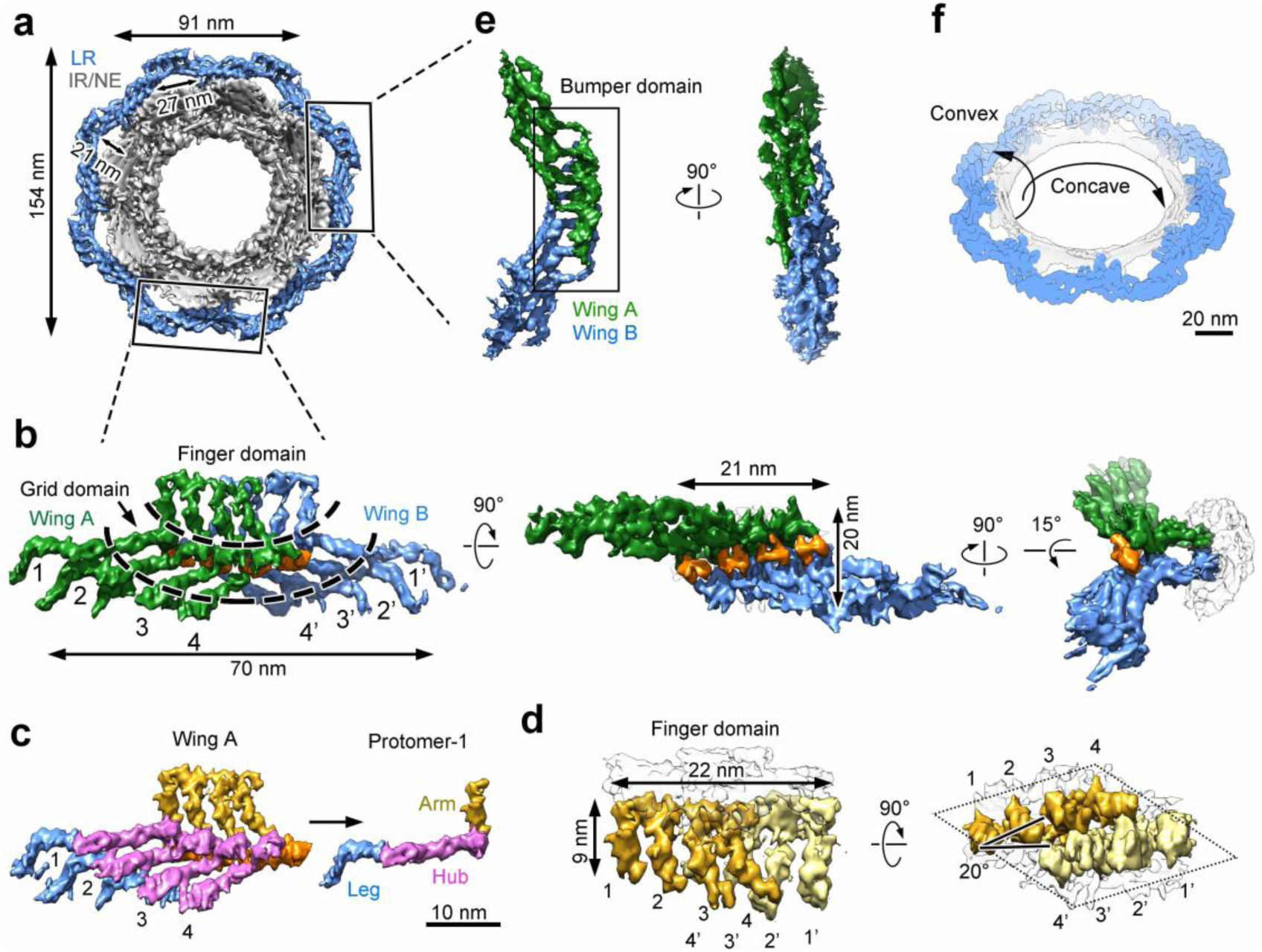
Structure of the LR subunit by the SPA approach. **a** Eight LR subunits form a continuous circular scaffold. The overall structure of the LR (colored marine) is viewed along the nucleocytoplasmic axis. **b** Structure of the LR subunit. Each butterfly-shaped LR subunit comprises two symmetric wings: Wing-A (colored green) and Wing-B (blue), which interact with each other through an extended interface (orange). The LR subunit has a Finger domain that contacts the fusion and a Grid domain that make up the bulk of the circular LR scaffold. Three mutually perpendicular views are shown. **c** Each wing of the LR subunit comprises four elongated, tubular protomers. These protomers (numbered 1 through 4) associate with each other in a planar fashion. Each protomer contains an arm (colored gold) at one end, a central hub (magenta), and a leg (blue) at the other end. Scale bar, 10 nm. **d** The Finger domain directly contacts the fusion of nuclear membranes. The tips of the protomers likely traverse the pore membrane and anchor the LR subunit to the pore. The cross section of the Finger domain has the shape of a diamond as indicated by dotted lines (right panel). **e** The Bumper domain is formed between two neighbouring LR subunits. Four legs from Wing-A of an LR subunit interact with four legs from Wing-B of the neighbouring LR subunit to form the Bumper domain. Two perpendicular views are shown. **f** The LR may stabilize both the concave and the convex curvatures of the fusion. The concave curvature relates to the diameter of the fusion within the NE, whereas the convex curvature is defined by the separation of the INM and ONM. Scale bar, 20 nm.

The Finger domain directly contacts the fusion between the INM and ONM (Fig. 3d and Supplementary information, Fig. S8d). The four arms 1-2-3-4 from one wing (or 1’-2’-3’-4’ from the other wing) constitute two pairs 1-2 and 3-4 (or 1’-2’ and 3’-4’ for the other wing), with the tips of each pair connected to each other (Fig. 3d, left panel and Supplementary information, Fig. S8d, left panel). Within the same Finger domain, three arms from one wing pair up with three arms from the other wing in a reciprocal order: 2-4’, 3-3’, and 4-2’. Together, the eight arms display a diamond-shaped cross section, with four arms from each wing constituting one side of the diamond (Fig. 3d, right panel and Supplementary information, Fig. S8d, right panel). The two distal arms, one from each wing (1 and 1’), are placed at the opposing ends of the diamond.

Two neighbouring LR subunits interact with each other through their legs, producing a characteristic scaffold that is hereafter referred to as the Bumper domain (Fig. 3e and Supplementary information, Fig. S8e). In contrast to the Finger domain, the Bumper domain is distal to the fusion and appears to cushion neighbouring NPCs (Fig. 1b and Supplementary information, Fig. S4). Both the Finger domain and the Bumper domain are visible in the original tomograms (Fig. 1a, arrowheads). Eight Grid domains and eight Bumper domains of the LR alternate to assemble into a closed ring scaffold, which places eight Finger domains in close contact with the fusion (Fig. 3a and Supplementary information, Fig. S8a). On one hand, the LR scaffold may stabilize the concave curvature of the fusion, which is defined by the diameter of the fusion within the equatorial plane of the NE (Fig. 3f and Supplementary information, Fig. S8f). On the other hand, the Finger domain directly contacts the luminal side of the fusion and may stabilize the convex curvature that connects INM to ONM. The spatial separation between INM and ONM is thought to be 10–30 nm^35^. The thickness of the LR subunit perpendicular to the nuclear membrane is approximately 20 nm (Supplementary information, Fig. S8b, right panel), which may help define the thickness of the NE surrounding the NPC.

### The Bumper domain of the LR

Back-projection of the LR subunit reconstruction into the original tomogram reveals two distinct lengths of the Bumper domains: 29 and 34 nm (Fig. 4a). The 29-nm Bumper domain has two legs from one wing pairing up with two legs from the other wing, producing six apparent legs as viewed along the nucleocytoplasmic axis (Fig. 4a). For this reason, the 29-nm Bumper domain is hereafter named Bumper-6. Similarly, the 34-nm Bumper domain is named Bumper-7 because the interface only involves one leg from each wing, leading to seven apparent legs (Fig. 4a). To reveal additional features, we identified and subjected 33,692 candidate Bumper domains to 3D classification with application of a local mask by the STA approach (Fig. 4b). A relatively large class number of 10 was applied to ensure identification of different conformations of the Bumper domain. Approximately 21.5% and 38.4% of all these domains, representing eight classes, belong to Bumper-6 and Bumper-7, respectively (Fig. 4b). The remaining two classes (39.4% in total) cannot be identified and likely represent deformed or damaged Bumper domains, or random noise.

**Fig. 4.**
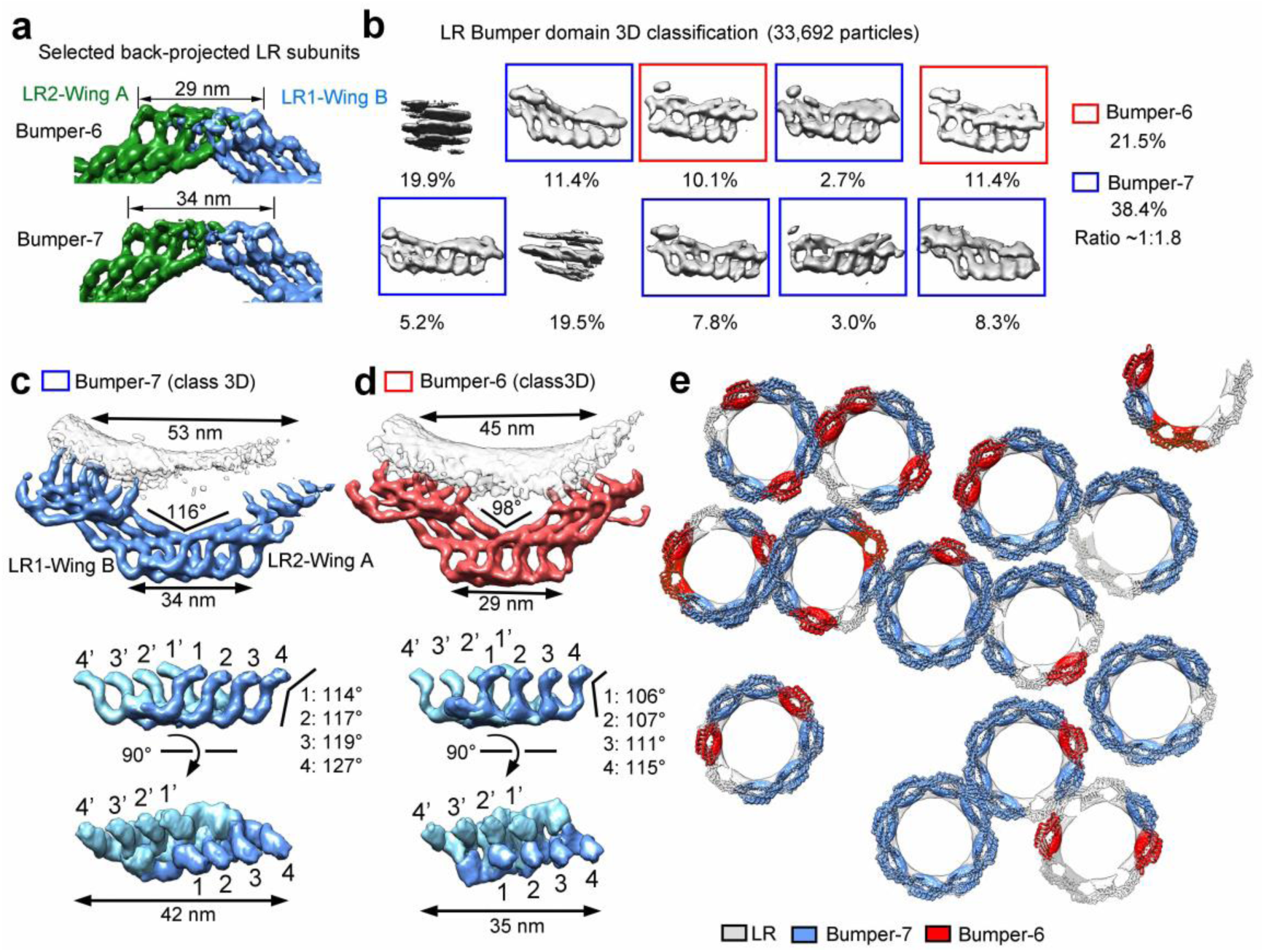
Structure of the Bumper domain of the LR by the STA approach. **a** The Bumper domain adopts two major conformations. By back-projecting the NPC particles onto the original tomograms, two major conformations of the Bumper domain are seen and named Bumper-6 and −7. The lengths of Bumper-6 and −7 are 29 and 34 nm, respectively. **b** Classification of the Bumper domains. The Bumper domains were re-cropped and classified. Bumper-6 and −7 represent 21.5% and 38.4% of the total particles classified. **c** Refined structure of Bumper-7 exhibits distinguishing features (upper panel). Two mutually perpendicular views are shown for an isolated Bumper-7 (middle and lower panels). **d** Refined structure of Bumper-6. In contrast to Bumper-7, the internal interface of Bumper-6 involves two pairs of promoter tips. **e** Mapping the Bumper domains onto the NPC particles. The reconstructions for Bumper-7 (marine), Bumper-6 (red), and the LR subunit (grey) were back-projected onto the original tomograms based on the refined coordinates of the individual particles. Shown here is a representative section of the nuclear membrane. The conformations of nearly all the classified Bumper domains are the same as those of the Bumper domains formed by independently back-projecting the LR subunits onto the original tomograms.

Additional refinement of the particles that belong to Bumper-6 and Bumper-7 reveals detailed features (Fig. 4c,d and Supplementary information, Fig. S9a,b). In Bumper-7, the first leg (leg-1) from one subunit pairs up with its corresponding leg (leg-1’) from an adjacent subunit (Fig. 4c). In Bumper-6, two legs from one subunit form two reciprocal pairs with two legs from an adjacent subunit: 1-2’ and 2-1’ (Fig. 4d). As a consequence of the different pairing arrangements, the centres of the two LR subunits that form Bumper-7 are separated by 53 nm (Fig. 4c), longer than the distance of 45 nm for Bumper-6 (Fig. 4d). The pairing difference also generates contrasting features in the overall structure as well as the angles extended between the hub and leg of the corresponding protomers (Fig. 4c,d).

These refined Bumper domains were projected back onto the original tomograms (Fig. 4e and Supplementary information, Fig. S9c,d). Bumper-6 and Bumper-7 are often seen in the same NPC, with Bumper-7 usually representing the majority. We speculate that the distinct Bumper conformations may reflect consequences of mechanical stress or deformation of the nuclear pore. In fact, most NPCs display a slightly elliptical appearance that amounts to a small change of the pore diameter^12^. Under these circumstances, the LR may help maintain the curvatures of the fusion through a conformational switch between Bumper-6 and Bumper-7.

### The Bumper domain cushions neighbouring NPCs

At least two factors – the extraordinarily large molecular mass and the relatively loose association among the four rings (CR/IR/NR/LR) – may make the NPC susceptible to mechanical deformation. There are approximately 2000-5000 NPCs in a vertebrate nucleus, with a density of 10-20 NPCs per square µm^9,36^. In contrast, a single *X. laevis* oocyte contains about 5×10^7^ NPCs, with a density of 60 NPCs per square µm^9,36^. In our STA reconstruction, NPC particles contact each other through their respective LRs (Figs. 1, 4e and Supplementary information, Figs S4, S9c-d). Judging from the cross section within the equatorial plane, a sizable fraction of the NPCs has been deformed into elliptical appearances. At every point of contact, the Bumper domains from one NPC cushion against the Bumper domains from neighbouring NPCs.

We performed a statistical analysis on deformation of the NPCs. Based on our reconstruction, a representative NPC has an outer diameter of 154 nm, with a distance of 130 nm between two Grid domains on the opposing sides of the LR (Fig. 5a, inset). An important judgment for deformation is whether the distance between the centres of two neighbouring NPCs is shorter than the standard outer diameter of an undeformed NPC. For each NPC, the distance to its closest neighbour is measured and plotted (Fig. 5a). Much to our surprise, the most frequently observed distance is 127.5 nm, which occurs to 1208 distinct pairs of NPCs. The average shortest distance between two neighbouring NPCs is 136 nm, which is 18 nm shorter than the outer diameter of a perfectly symmetric NPC. This analysis suggests widespread deformation or crowding of the NPCs on the *X. laevis* oocyte. Our observed distance is in excellent agreement with the reported average pairwise distance of 135±5 nm for the NPCs derived from stage VI *X. laevis* oocytes^37^.

**Fig. 5.**
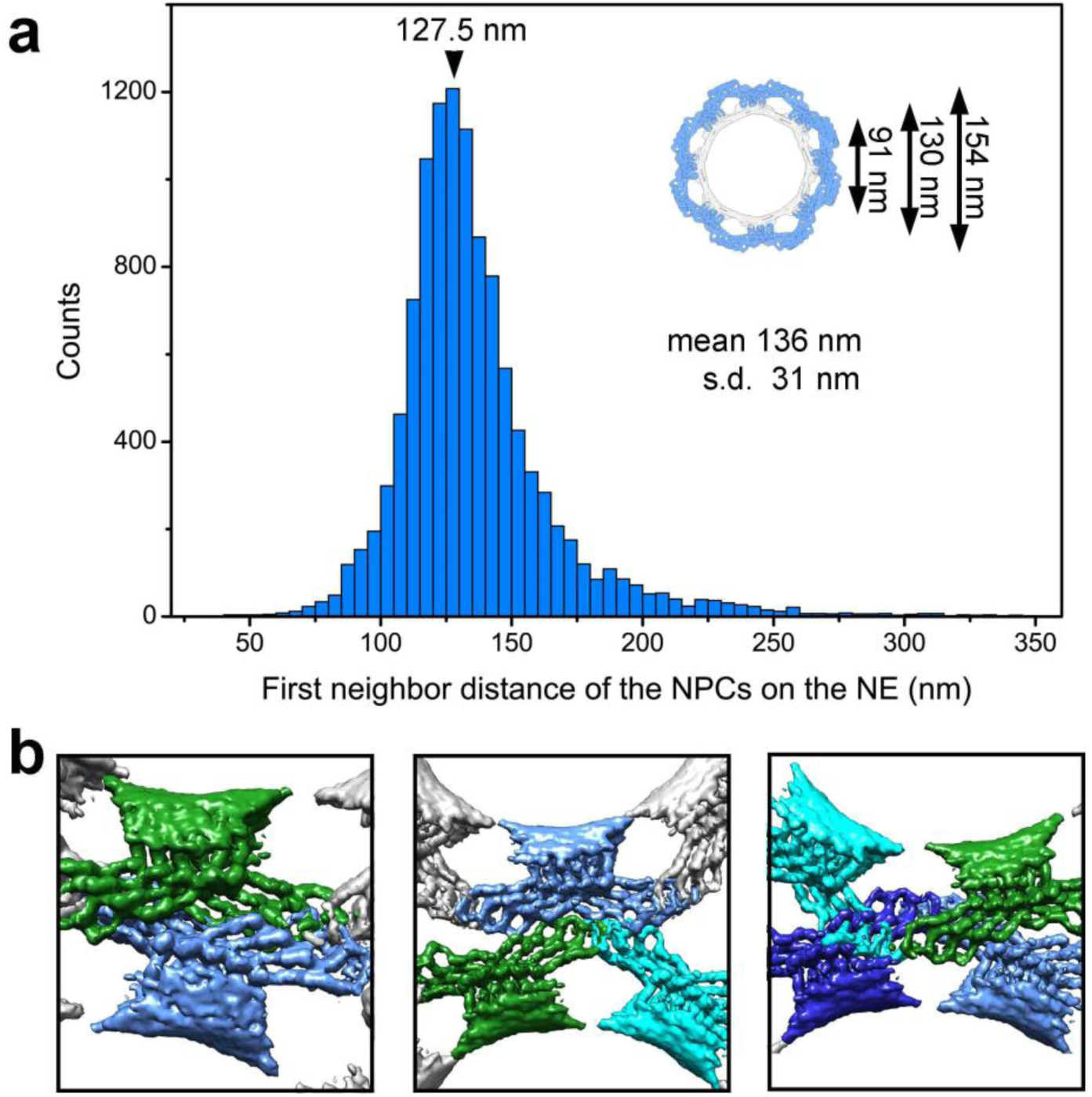
The Bumper domain may cushion neighbouring NPCs. **a** NPCs are generally deformed in the *X. laevis* nuclear membrane. Statistical analysis of the deformation among neighbouring NPCs. For each NPC, the distance to its nearest neighbor is measured and plotted here. The median distance is 136±31 nm. The most frequently observed distance is 127.5 nm, which occurs to 1208 pairs of NPC. The sizes of a symmetric LR is shown in the inset for reference. **b** Three representative examples of close contact between neighbouring NPCs. The reconstruction for the LR subunit was back-projected onto the original tomograms based on the refined coordinates of the individual LR subunit particles.

Close examination of the tomograms reveals striking examples of crowding among neighbouring NPCs (Fig. 5b). The Bumper domain of one NPC clashes with the Bumper domain of a neighbouring NPC (Fig. 5b, left panel) or invades into the space between two Bumper domains of a neighbouring NPC (Fig. 5b, middle panel). Under extreme circumstances, the Bumper domain of one NPC slides past the Bumper domain of a neighbouring NPC, allowing the Grid domains to contact each other (Fig. 5b, right panel). Notably, among all NPCs examined, no Bumper domain reaches the membrane fusion of a neighbouring NPC.

## DISCUSSION

Structures of the NPC have been reported for at least five organisms^7,12-15^ (Supplementary information, Fig. S7). In this study, we report the cryo-ET structure of the NPC from *X. laevis* oocyte (Supplementary information, Fig. S3), which allows identification of previously unknown features of the LR (Figs. 2a, 4 and Supplementary information, Fig. S8). Preferred orientation of the NPC particles led to resolution anisotropy, particularly along the Z-axis (Supplementary information, Fig. S3c,d), which may have an impact on the cylindrical height of the NPC. Nonetheless, such anisotropy has little impact on the general conclusions derived from the STA-based cryo-ET study, in part because the key features of the LR subunit have been validated by an independent SPA-based cryo-EM study (Figs. 2b, 3). In the SPA-based reconstruction, the edge of the two wings in the LR subunit is less well defined compared to the STA-based reconstruction (Fig. 2c); this is due to application of a considerably smaller alignment mask for SPA compared to STA. Despite differences in the mask size, all eight legs in the LR subunit are clearly visible in both approaches (Supplementary information, Fig. S10a,b). Importantly, reconstruction by SPA agrees with that by STA up to 13.2 Å using a common mask for the FSC (Supplementary information, Fig. S10c). The observed structural features of the LR subunit are previously unknown and appear to define and perhaps explain the function of the LR (Figs. 3-5).

Despite its mysterious nature, existence of the LR has been previously recognized. The NPC from *Necturus maculosus* and *X. laevis* was observed to contain eight spokes that penetrate the fusion into the lumen and form a “luminal ring” through radial arm dimers or handle-like luminal domain^14,16-19,38^. In contrast, the luminal structure in the *D. discoideum* NPC appears to comprise eight discrete rods^12^. *H. sapiens* NPC also contains luminal connections^20^. GP210 was found to form a ring around the NPC from *X. laevis*^29^. The EM structure of Pom152 has an extended, tubular appearance^39,40^. Reconstruction of the *S. cerevisiae* NPC shows a fusion-associated ring scaffold with eight arches, each speculated to comprise two Pom152 molecules^7,40^. These previous studies mostly report rough overall appearance of the LR scaffold. In this study, we have identified key structural features of the LR from *X. laevis* NPC using both cryo-ET and cryo-EM.

The lack of structural information on any of the candidate Nups of the LR, together with the limited resolution of our EM reconstruction, do not allow conclusive identification of the protein components in the density of the LR subunit. Nonetheless, among the candidate components of the LR, only GP210 contains a luminal domain that is large enough to constitute the LR scaffold observed in our study^17,28^. *X. laevis* GP210 contains 1,898 residues and was predicted to contain 15 immunoglobulin-like (Ig-like) domains^39^ (Supplementary information, Fig. S11a). It is likely that GP210 constitutes the bulk of the LR protomer. Supporting this analysis, GP210 was found to form a ring around the *X. laevis* NPC, with an eight-fold symmetry and a diameter of 164±7 nm^29^. In contrast to GP210, POM121 only has its N-terminal ∼30 residues in the lumen and NDC1 only has the linker sequences between neighbouring TMs in the lumen^4^ (Supplementary information, Fig. S11b). Therefore, the bulk of the elongated tubular density of the LR protomer may come from the 15 Ig-like domains of GP210 (Supplementary information, Fig. S11c). Despite these tantalizing clues, we cannot exclude the possibility that the LR is formed by other yet-to-be identified proteins.

Few *X. laevis* NPC particles display a perfectly eight-fold symmetry, and most are slightly elliptical in appearance (Figs 1, 4e and Supplementary information, Fig. S9c,d). The shape asymmetry could arise as a result of osmotic swelling or temperature variation^17,18^. Although we cannot exclude the possibility of NPC deformation during sample preparation, examination of the *D. discoideum* NPC on intact nuclei and *H. sapiens* NPC in whole cells also revealed radial displacement and elliptical distortions^12,20^. Asymmetric variations and diameter dilation at the level of individual NPCs were also observed in HeLa cells^41^, and the algae NPC displayed a dilated IR^13^; in both cases, the samples were prepared through focused ion beam (FIB) milling that was supposed to maintain the original appearance in cells. Taken together, the NPC may adopt a much more dynamic conformation than anticipated^13^. The observed deformation of the NPC may reflect its natural state on the NE and such plasticity could be indispensable to its biological functions^18^.

Assuming GP210 is the primary constituent of the LR scaffold, our finding that the Bumper domain cushions neighbouring NPCs may mechanistically explain the observation that GP210 mediates nuclear pore formation, dilation, NPC spacing and integrity^42,43^. Knock-down of GP210 in HeLa cells and *Caenorhabditis elegans* led to clustering of NPCs in dying cells^43^, likely due to loss of such cushioning. The Bumper domain appears to exhibit marked elasticity, with Bumper-7 being the default state. Any force squeezing the NPC towards the centre may cause a group of four protomers within one wing to slide towards that within another wing of an adjacent LR subunit, thus switching Bumper-7 to Bumper-6 (Fig. 4c,d). This speculated property of the NPC may serve to absorb radial shock and insulate the transport function from movements of the nuclear membrane.

Two wings from adjacent LR subunits constitute an arch, reminiscent of the LR in yeast^7,40^. Relative to the pore membrane, each arch defines a passage that measures 27 nm in width (Fig. 3a and Supplementary information, Fig. S8a). As noted previously^7,40^, these passages are aligned with the circumferential passages between the CR or NR and the pore membrane, and between neighbouring IR subunits. Together, these passages could form lateral openings between subunits of the NPC. The combination of arches and passages may outline a well-defined duct for nucleocytoplasmic transport of INM proteins^44,45^.

Cryo-ET reconstruction also allows visualization of other macromolecular complexes on the tomograms. Analysis of 14 tomograms allowed preliminary reconstruction of the ribosomal subunits, translocon-associated protein complex (TRAP) and oligosaccharyl transferase (OST)^46^ (Fig. 1 and Supplementary information, Fig. S4). As observed in a previous study^41^, such complexes are present in a large number on the cytoplasmic side of the ONM. In-depth examination of these tomograms may reveal additional molecular machineries that associate with the NPC.

## ACKNOWLEDGEMENTS

We thank Westlake University for providing a Start-up fund, the Tsinghua University Branch of China National Center for Protein Sciences (Beijing) for the cryo-EM facility and the computational facility support, and L. Zhao, X. Li, and J. Wen for technical support. We thank X. Fu and P. Zhang at the University of Pittsburgh for advice on STA sample preparation and SerialEM data collection. This work was supported by funds from the National Natural Science Foundation of China (31930059, 81920108015, 31621092 and 31430020).

## AUTHOR CONTRIBUTIONS

X.Z. and Y.Z. prepared the sample. Y.Z., C.Z., G.H., S.L., Q.W. and J.L. collected the EM data. Y.Z., S.L., G.H., C.Z. and Q.W. processed the EM data. Y.Z. and S.L. performed the cryo-ET STA calculation. G.H. performed the cryo-EM SPA calculation. K.W., W.Z. and G.Y. provided computing assistance. Q.Z., C.Y. and Q.T. provided critical advices. All authors analyzed the structure. Y.Z., S.L., G.H., C.Z. and Y.S. wrote the manuscript. Y.S. conceived and supervised the project.

### Competing interests

The authors declare no competing financial interests. Correspondence and requests for materials should be addressed to Y.S.

## MATERIALS AND METHODS

### Cryo-sample preparation

Small pieces of ovary were separated from a 2-4 years old, narcotized female frog (Nasco, USA) and transferred into modified Barth’s saline (MBS) (10 mM HEPES, pH 7.5, 88 mM NaCl, 1 mM KCl, 0.82 mM MgSO_4_, 0.33 mM Ca(NO_3_)_2_ and 0.41 mM CaCl_2_) at 18°C. Stage VI oocytes (a sphere of 1.3 mm in diameter with clear separation of the black animal pole, the off-white vegetal pole and unpigmented equatorial belt) were manually sorted from the encased connective tissue membranes using forceps. For each stage-VI oocyte, a small hole was generated on the side of the animal pole using forceps in MBS, and the nucleus was sucked and transferred into a low salt buffer (LSB) (10 mM HEPES, pH 7.5, 1 mM KCl, 0.5 mM MgCl_2_, 10 μg/ml aprotinin, 5 μg/ml leupeptin) using a 20-μl pipette tip. The isolated nuclei were kept in LSB for 10 minutes, and during this time the yolk was rapidly cleaned through pipetting several times^14,47^. Two-to-three cleaned nuclei were transferred onto a freshly glow-discharged copper EM grid (R1.2/1.3; Quantifoil, Jena, Germany) in an LSB liquid drop of about 5-μl.

The glow-discharged grids were prepared for 30 seconds using the “Mid” setting of the Plasma Cleaner (Harrick, Plasma Cleaner PDC-32G). For each nucleus, the nuclear envelope (NE) was spread onto the EM grid by popping a small hole on one side of the nucleus using two glass needles to extrude chromatin and other nuclear contents. The NE was carefully washed three times, each time using 5 μl LSB. 2 μl gold fiducial beads (10 nm diameter, Aurion, The Netherlands) were applied onto the native NE samples before plunge-freezing. The grids were blotted for 5 seconds, vitrified by plunge-freezing into liquid ethane using a Vitrobot Mark IV (Thermal Fisher Scientific) at a temperature of 8°C and a humidity of 100%. The quality of sample preparation was examined using an FEI Tecnai Arctica microscope (Thermo Fisher Scientific) operating at 200 kV.

### Cryo-ET data acquisition

The grids were imaged on a Titan Krios microscope operating at 300 kV equipped with an energy filter (slit width 20 eV; GIF Quantum LS, Gatan) and a K2 Summit direct electron detector (Gatan). Tilt-series were recorded in the super-resolution mode at a nominal magnification of 64,000x, resulting in a calibrated pixel size of 1.111 Å. A combination of dose-symmetric^30^, bi-directional, and continuous schemes were used to collect tilt-series from −60° to 60° at a step size of 3° using SerialEM^31^ (Supplementary information, Fig. S1). At each tilt, a movie stack consisting of 8 frames was recorded at 0.1 second exposure per frame, yielding a total dose of ∼90-150 e^-^/Å^2^ per tilt-series. 1,575 tilt-series were collected using defocus values between −2 and −4 µm (Supplementary information, Fig. S2). A summary of data acquisition statistics can be found in Supplementary information Table S1.

### Cryo-ET data processing

Tilt-series were binned to a final pixel size of 2.22 Å and motion corrected by averaging eight frames for each tilt using MotionCor2 (ref. 48). Defocus of the tilt series was estimated using CTFFIND4 (ref. 49). The contrast transfer function (CTF) of each tilt-series was corrected using NovaCTF^50^. 1,425 tilt-series with good fiducial alignment were reconstructed to tomograms through weighted back projection using IMOD^51^ (Supplementary information, Fig. S2). The tomograms were 2× and 4× binned for subsequent processing. Subtomograms of 36,529 NPC complexes were manually picked and extracted from the 4× binned tomograms into boxes of 200×200×200 voxels with the help of Dynamo catalogue^52^ for further analysis.

Sub-tomogram averaging (STA) was carried out in Dynamo^32^, following a published protocol^53^. For reconstruction of the cytoplasmic ring (CR), the manually picked 4× binned NPC sub-tomograms were averaged as the template for their alignment. The resolution was restricted to 40 Å and a C8 symmetry was applied at this stage. Next, coordinates of 292,232 CR subunits were estimated by subboxing and the subboxes were extracted from the 4× binned tomograms into boxes of 100×100×100 voxels for independent alignment. To prevent overfitting, a ‘gold-standard’ method using an adaptive filter and an ellipsoidal mask was used to align the subboxes. The aligned CR subunits were subjected to 3D classification (particles binned to 8×) using RELION3.0 (ref. 33). 112,220 subunits survived this analysis and were further aligned in Dynamo to 17.8 Å resolution using a criterion of 0.143 for the Fourier shell correlation (FSC) value. Next, the refined coordinates were used to extract and align subtomograms from 2× binned tomograms. Finally, subtomograms from the unbinned tomograms were extracted into boxes of 320×320×320 voxels and aligned to a final resolution of 9.1 Å.

For reconstruction of the inner ring (IR), the 112,220 4× binned CR subtomograms were aligned using the human IR subunit density map (EMD-3106)^15^ as the initial template and an ellipsoid at the IR location as the mask (Supplementary information, Fig. S2). After aligning the IR subunits to 17.8 Å resolution, the boxes were re-centered to the IR subunit and the coordinates were used to extract 2× binned IR subtomograms into boxes of 200×200×200 voxels. 3D classification by RELION3.0 was used to select a group of 34,086 IR subunits. Subsequent alignment in Dynamo achieved an average resolution of 13.1 Å for the IR.

For reconstruction of the nuclear ring (NR), the 112,014 4× binned and re-centered IR subtomograms were aligned using human NR subunit density map (accession code EMD-3107)^15^ as the initial template and an ellipsoid at the NR location as the mask (Supplementary information, Fig. S2). The NR subunit density emerged after initial alignment, which was restricted at 60 Å resolution to prevent template bias. Subsequently the boxes were re-centered to the NR subunit and further aligned to 17.8 Å resolution. 3D classification by RELION3.0 was used to select a group of 34,068 NR subunits, which were extracted into 2× binned NR subtomograms with a box size of 200×200×200 voxels. Subsequent alignment achieved an average resolution of 13.6 Å for the NR.

For reconstruction of the luminal ring (LR), the 112,014 4× binned and re-centered IR subtomograms were aligned using the aligned IR subunits as initial templates and an ellipsoid at the LR location as the mask (Supplementary information, Fig. S2). The LR subunit density emerged after initial alignment. The boxes were re-centered to the LR subunit and further aligned to 17.8 Å resolution. 3D classification by RELION3.0 was used to select a group of 34,087 LR subunits, which were extracted into 2× binned LR subtomograms with a box size of 200×200×200 voxels. Subsequent alignment achieved an average resolution of 15.1 Å for the LR. To further examine the conformation of the Bumper domain, the boxes were re-centred to the joint region between two adjacent subunits of the LR. 3D classification of the subtomograms by RELION3.0 revealed two major conformations: Bumper-6 and Bumper-7. Using Dynamo, 7,277 Bumper-6 and 13,087 Bumper-7 particles were independently aligned and refined, yielding average resolutions of 17.0 Å and 15.6 Å, respectively (Supplementary information, Fig. S9a, b).

To illustrate the organization of the NE, ribosomes were reconstructed. 780 ribosomes were manually picked from 14 4×binned tomograms and extracted into boxes of 80×80×80 voxels. The sub-tomograms were first aligned using EMD-4315 (ref. 54) as the template, and the resolution was restricted to 45 Å at this stage. Next, the refined coordinates were used to extract subtomograms into boxes of 160×160×160 voxels from 2× binned tomograms, which were subsequently aligned using the FSC standard of 0.143 to 16.4 Å resolution. Consistent with reported ribosome reconstruction from the NE^41^, some of the ribosomes reconstructed here are also found to be associated with translocon-associated protein complex (TRAP) and oligosaccharyl transferase (OST)^46^.

The subunit maps were low-passed according to the estimated local resolutions of the reconstructions^55^. Empirical B-factors of −1000, −2000, −2000, −2000 and −2000 were used to sharpen the CR, IR, NR, LR and ribosome reconstructions, respectively.

### Cryo-EM data acquisition

Details for the acquisition of cryo-EM data are described in the accompanying manuscript^34^. Briefly, micrographs were recorded on a Titan Krios (FEI) electron microscope, operating at 300 kV and equipped with a Gatan Gif Quantum energy filter (slit width 20 eV). A K2 Summit detector (Gatan Company) in super-resolution mode with a nominal magnification of 64,000x was used, resulting in a calibrated pixel size of 1.111 Å. The total dose followed a cosine alpha scheme where the total dose is inversely proportional to the cosine of the tilting angle and the total dose used for the Tilt-0 micrographs was 52 e^-^/Å^2^, a summary of data acquisition statistics can be found in Supplementary information Table 2.

### Cryo-EM data analysis

The single-particle cryo-EM data was mainly used to generate a reconstruction of the CR at an improved resolution^34^. After completing this task, the same data was analyzed to generate a SPA-based reconstruction of the LR. The strategy for processing of the cryo-EM data towards reconstruction of the LR subunit is presented in Supplementary information, Fig. S5. We first attempted to reconstruct the LR subunit by simply re-centering the particles to the LR subunit and preformed image alignment with or without any mask. This approach however failed with the CR subunit showing strong density at the edge of the box or the mask that was supposed to emphasize the LR subunit. In order to resolve this problem, two measures were implemented.

First, since the whole image alignment was biased towards the CR subunit, we tested whether placing a CR subunit on the opposite side could somehow neutralize this bias. Results from the cryo-ET study indicates a two-fold symmetry of the LR subunit at up to 15-Å resolution. Specifically, we updated the Euler angle and offsets for each particle to generate another RELION data star file. This effort resulted in an identical reconstruction when the whole reconstruction was rotated by 180 degrees, so that another CR subunit would appear at the bottom of the box instead of the top (Supplementary information, Fig. S5). The particles within this star file are referred to as symmetry related particles, which were then joined by the original particles to generate a data set with twice the number of particles as the original data set. Second, the CR subunit appeared partially because the CTF parameter favored high resolution reconstructions of the CR subunit, this would likely bias the initial stages of alignment of the LR subunit towards the CR side because of its strong features. To resolve this potential problem, we removed all CTF parameters from previous CTF-refinements and reverted back to the CTF values from the micrograph star file.

These two strategies allowed us to generate an initial reconstruction of the LR subunit. This initial reconstruction was refined with application of a soft mask that covers the Finger domain, the Grid domain and the legs. To further improve the resolution, six rounds of 3D classifications were performed to remove empty or heterogeneous particles. This practice results in removal of 20 to 30 percent of the particles after each round of 3D classification. The final average resolution of the reconstruction for the LR subunit was 10.7 Å using 311,240 particles, which includes 157,541 particles from the original data set and 153,699 particles from the symmetry related data set.

### Data deposition

The Electron Microscopy Database (EMD) accession codes of the LR subunit, Bumper-7, Bumper-6, the CR subunit, the IR subunit and the NR subunit are EMD-0983, EMD-0984, EMD-0985, EMD-0986, EMD-0997 and EMD-0998, respectively, for the reconstructions calculated by the STA approach. The EMD accession code is EMD-0982 for the reconstruction of the LR subunit calculated by the SPA approach.

