## Supplementary information for "Molecular architecture of the luminal ring of the *Xenopus laevis* nuclear pore complex"

Short title: Structure of the NPC luminal ring

### Yanqing Zhang^1,2,6^, Sai Li^3,4,6^, Chao Zeng^4,6^, Gaoxingyu Huang^3,6^, Xuechen Zhu^1,2,6^, Qifan Wang^1,2^, Kunpeng Wang^5^, Qiang Zhou^1,2^, Wusheng Zhang^5^, Guangwen Yang^5^, Minhao Liu^3^, Qinghua Tao^3^, Jianlin Lei^3^, and Yigong Shi^1,2,3,4,*^

^1^Key Laboratory of Structural Biology of Zhejiang Province, School of Life Sciences, Westlake University, 18 Shilongshan Road, Hangzhou 310024, Zhejiang, China; ^2^Institute of Biology, Westlake Institute for Advanced Study, 18 Shilongshan Road, Hangzhou 310024, Zhejiang Province, China

^3^Beijing Advanced Innovation Center for Structural Biology & Frontier Research Center for Biological Structure, ^4^Tsinghua University-Peking University Joint Center for Life Sciences, School of Life Sciences, Tsinghua University, Beijing 100084, China; ^5^Tsinghua Computing Facility & Department of Computer Science, Tsinghua University, Beijing 100084, China

^6^These authors contributed equally to this work.

**SUPPLEMENTARY FIGURE LEGENDS**

**Supplementary information, Fig. S1 | Representative 0°-tilt micrographs and the associated defocus values for three tilt-schemes used in this study.** **a**, Dose-symmetric tilt-scheme (-60° to 60°). 240 tilt-series were recorded. Shown here (tomo 586) is a representative example. **b**, Bidirectional tilt-scheme (0° to -60° and 3° to 60°). 874 tilt-series were recorded. Shown here (tomo 453) is a representative example. **c**, Continuous tilt-scheme (-60° to 60°). 311 tilt-series were recorded. Shown here (tomo 687) is a representative example. In each of the three cases, the 0°-tilt micrograph and the defocus value for the specified tilt-series are shown in the upper and lower panels, respectively.

**Supplementary information, Fig. S2 | A processing flow chart of the cryo-ET data on the NPC from *Xenopus laevis* (*X. laevis*).** For detailed description, please refer to the Methods. CR, cytoplasmic ring; IR, inner ring; LR, luminal ring; NR, nuclear ring.

**Supplementary information, Fig. S3 | Average resolution and anisotropy of the cryo-ET reconstructions for the NPC from *X. laevis*.** **a**, The STA-based cryo-ET reconstructions of the CR, IR, NR, and LR subunits. The resolution range for each of these reconstructions is color-coded. **b**, The Fourier shell correlation (FSC) curves for the reconstructions of the CR, IR, NR, and LR subunits. On the basis of the FSC criterion of 0.143, the reconstructions of the CR, IR, NR, and LR subunits have average resolutions of 9.1 Å, 13.1 Å, 13.6 Å, and 15.1 Å, respectively. **c**, The cryo-ET reconstruction is anisotropic. Shown here are the Fourier shell correlation (FSC) curves of the CR, IR, NR and LR subunits in three different orientations. The FSC curves for tilt 0°, 45°, and 90° (named Tilt-0°, Tilt-45° and Tilt-90°) were calculated with cone masks in Fourier space and colored red, blue, and green, respectively. As a reference, the FSC curve for the global CR subunit is shown and colored black. The resolutions were calculated on the basis of the FSC criterion of 0.143. **d**, Angular distribution of particles for reconstruction of the CR, IR, NR and LR subunits.

**Supplementary information, Fig. S4 | Three-dimensional organization of the NPC particles in a local region of the *X. laevis* NE.** Four consecutive tomographic 2D slices are displayed in the left columns of panels **a** through **d**, with panel **a** on the cytoplasmic side and panel **d** on the nucleoplasmic side. These slices, each measuring 8.89 Å in thickness, are evenly spaced with a separation of 17.78 nm between two neighbouring slices. Structural interpretation of the regions is shown in the right columns of panels **a** through **d**, where the reconstructions for the NPC subunits and ribosomes were back-projected onto the original tomograms based on the refined coordinates of the individual particles. Ribosomes: 40S: Small ribosome subunit; 60S: Large ribosome subunit; TRAP: translocon-associated protein complex; OST: oligosaccharyl transferase.

**Supplementary information, Fig. S5 | A flow chart for the processing of the cryo-EM data on the LR subunit of the NPC from *X. laevis*.** Single-particle analysis (SPA) was applied in data processing. For detailed description, please refer to the Methods.

**Supplementary information, Fig. S6 | Reconstruction of the LR subunit of the *X. laevis* NPC by the SPA-based cryo-EM approach.**  **a**, The SPA cryo-EM reconstruction of the LR subunit. The resolution range is color-coded. Two mutually perpendicular views are shown. The corresponding cylinder representations of angular distribution are shown on the right. **b**, The FSC curve for the SPA-based reconstruction of the LR subunit. On the basis of the FSC criterion of 0.143, the reconstruction of the LR subunit has an average resolution of 10.7 Å.

**Supplementary information, Fig. S7 | Structural comparison of the cryo-ET reconstructions of the NPC.** Surface representations of cryo-ET reconstructions are shown to scale for *X. laevis* (this study), *X. laevis* (accession code EMD 3005-3008)^14^, *Homo sapiens* (EMD 3103)^15^, *Chlamydomonas reinhardtii* (EMD 4355)^13^ and *Saccharomyces cerevisiae* (EMD 7321)^7^. Three views of each reconstruction are shown: cross-section side view, 45°-tilt view, and top view from cytoplasmic side. The LR observed in this study is highlighted in marine.

**Supplementary information, Fig. S8 | Structure of the LR subunit by the STA approach. a**, Eight LR subunits form a circular scaffold around the fusion between the inner (INM) and outer nuclear membranes (ONM). The NPC is viewed along the nucleocytoplasmic axis. Two complete LR subunits are highlighted. Within each LR subunit, two wings are colored green and marine. **b**, Structure of the LR subunit. Each butterfly-shaped LR subunit comprises two symmetric wings: Wing-A (green) and Wing-B (blue), which interact with each other through an extended interface (orange). The LR subunit has a Finger domain and a Grid domain. Three mutually perpendicular views are shown. **c**, Each wing of the LR subunit comprises four planar, elongated, tubular protomers (numbered 1 through 4). Each protomer contains an arm (colored gold), a hub (magenta), and a leg (blue). The four protomers in each wing form two pairs (1-2 and 3-4), with the arms of each pair connected to each other through their tips. Scale bar, 10 nm. **d**, The Finger domain directly contacts the fusion of nuclear membranes. **e**, The Bumper domain is formed between two meighboring LR subunits. Four legs from Wing-A of an LR subunit interact with four legs form Wing-B of the neighboring LR subunit to form the Bumper domain. **f**, The LR may stabilize both the concave and the convex curvatures of the fusion. The Finger domain directly contacts the fusion and may stabilize the convex curvature. The LR surrounds the fusion in the plane of the NE and may strabilize the concave curvature. Scale bar, 20 nm.

**Supplementary information, Fig. S9 | Cryo-ET analysis of Bumper-6 and Bumper-7.** **a**, The local resolutions for Bumper-6 and Bumper-7 are color-coded in the cryo-ET reconstructions. Scale bar, 10 nm. **b**, The FSC curves for Bumper-6 (red) and Bumper-7 (blue). The resolutions for Bumper-6 and -7 are estimated to be 17.0 Å and 15.6 Å, respectively, on the basis of the FSC criterion of 0.143. **c**, A local region of the tomogram with the refined NPC particles. The reconstructions for Bumper-7 (marine), Bumper-6 (red), and the LR subunit (grey) were back-projected onto the original tomograms based on the refined coordinates of the individual particles. **d**, Another local region of the tomogram with the refined NPC particles. Coloring scheme is the same as in panel **c**.

**Supplementary information, Fig. S10 | Comparison of the STA and SPA reconstruction of the LR subunit. a**, A central slice of the LR reconstruction by the STA approach. The average local resolution in the Finger and Grid domains is 12.6 Å. **b**, A central slice of the LR reconstruction by the SPA approach. For both the STA and SPA reconstructions, similar features are observed for the Finger and the Grid domains. In contrast to STA-based reconstruction, the two Bumper domains are missing in the SPA-based reconstruction, due to application of a considerably smaller alignment mask. **c**, The FSC curve of the SPA reconstruction with the STA reconstruction shows agreement of up to 13.2 Å based on the FSC criterion of 0.143. A common mask was used.

**Supplementary information, Fig. S11 | Domain structure of four candidate integral membrane proteins in the LR.** **a**, A schematic diagram of the four candidate protein components of the vertebrate LR. The four proteins and the domains within each protein are drawn to scale. GP210 (Pom152 in yeast) is the largest of the three proteins, with 1898 amino acids. GP210 sequentially contains 15 predicted Ig-like domain, a C-domain rich in β-strands, a TM, and about 57 residues at the C-terminus. **b**, A topology diagram of the four candidate protein components of the LR. For GP210, all sequences N-terminal to the TM are in the lumen of the NE. In contrast, for NDC1, only the sequences connecting the six TMs are in the lumen; for POM121, only the N-terminal 31 residues reside in the lumen. TMEM33 has six predicted TMs and is the vertebrate homologue of yeast Pom33 but has not yet been experimentally confirmed as a Nup. **c**, The cryo-ET density of Pom152 (the yeast homolog of GP210), colored yellow here, is placed into the reconstruction for one LR protomer (colored grey). Pom152 is thought to contain nine Ig-like domains^40^.

**Supplementary information, Table S1 | Cryo-ET data collection and STA reconstruction statistics.**

**Supplementary information, Table S2 | Cryo-EM data collection and SPA reconstruction statistics.**

**Supplementary information, Video S1 | Visualization of macromolecular complexes across the NE on a representative tomogram.** Reconstructions of the CR, IR, NR, LR and ribosomal subunits were individually projected back into a representative tomogram (low-passed to 80 Å) containing a piece of 824×824 nm nuclear envelope, which is viewed from three distinct angles. Ribosomes: 40S: Small ribosome subunit; 60S: Large ribosome subunit; TRAP: translocon-associated protein complex; OST: oligosaccharyl transferase.
