## Supplementary figures and images for "Molecular architecture of the luminal ring of the *Xenopus laevis* nuclear pore complex"

### Supplementary information, Fig. S1

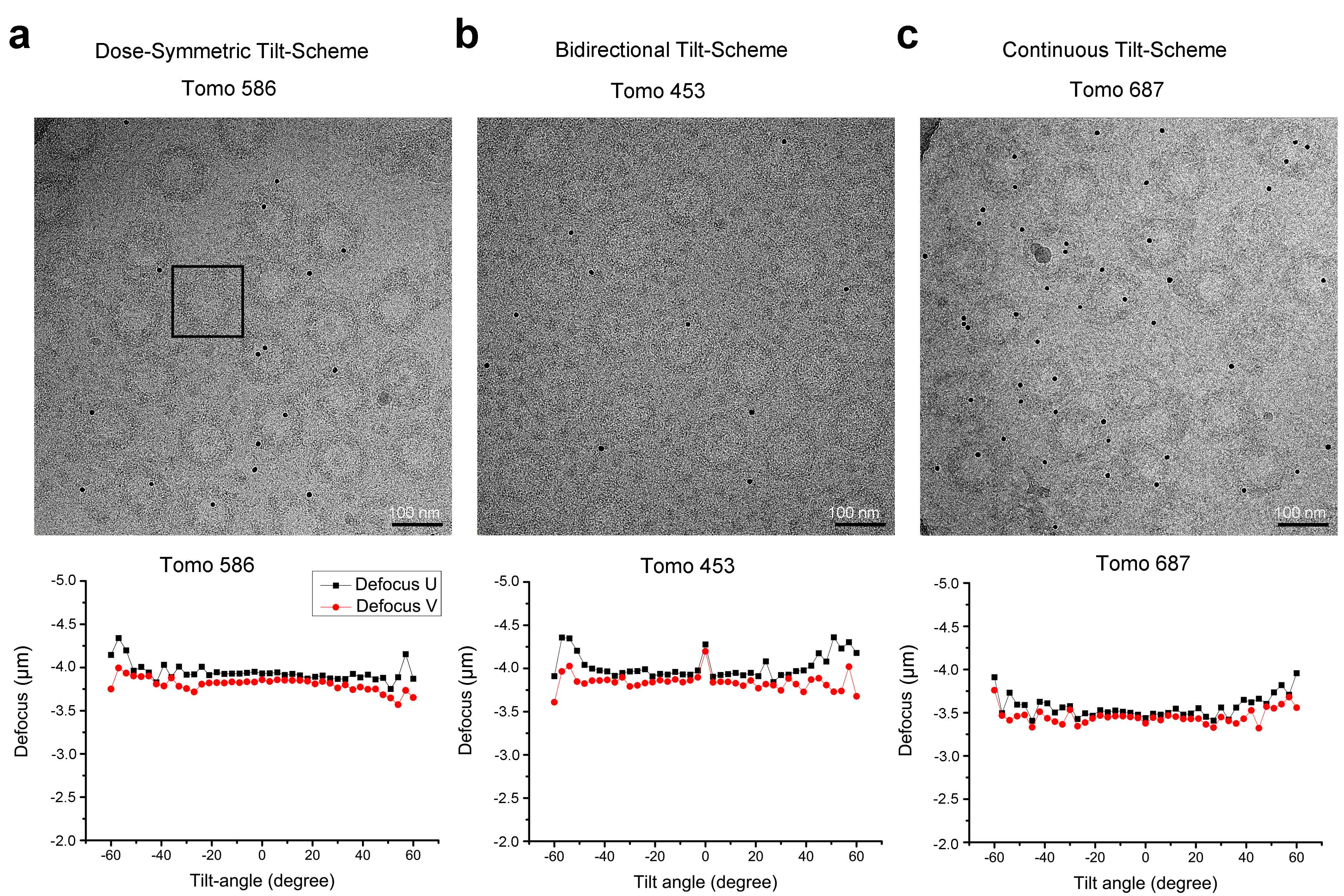

### Supplementary information, Fig. S2

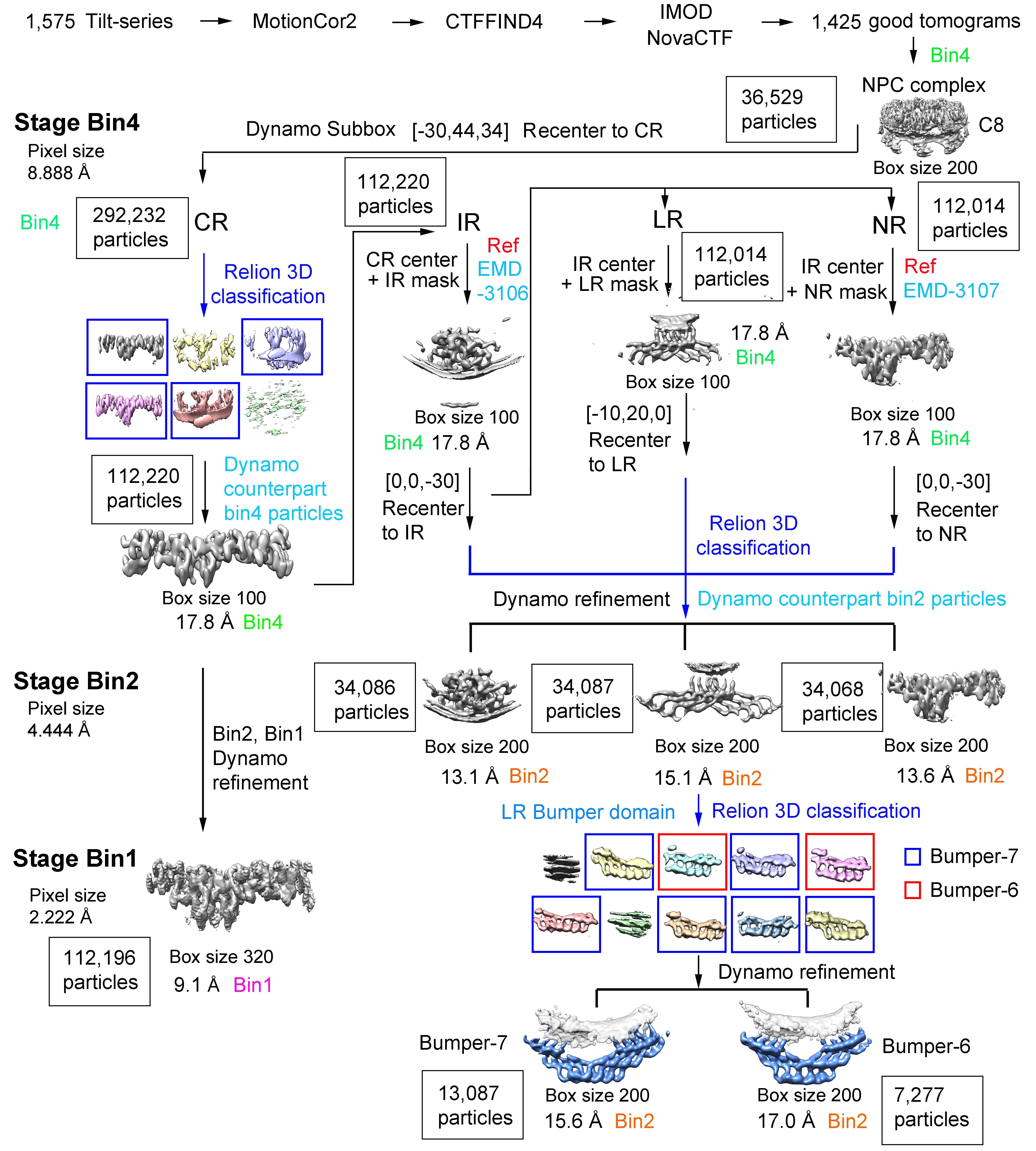

### Supplementary information, Fig. S3

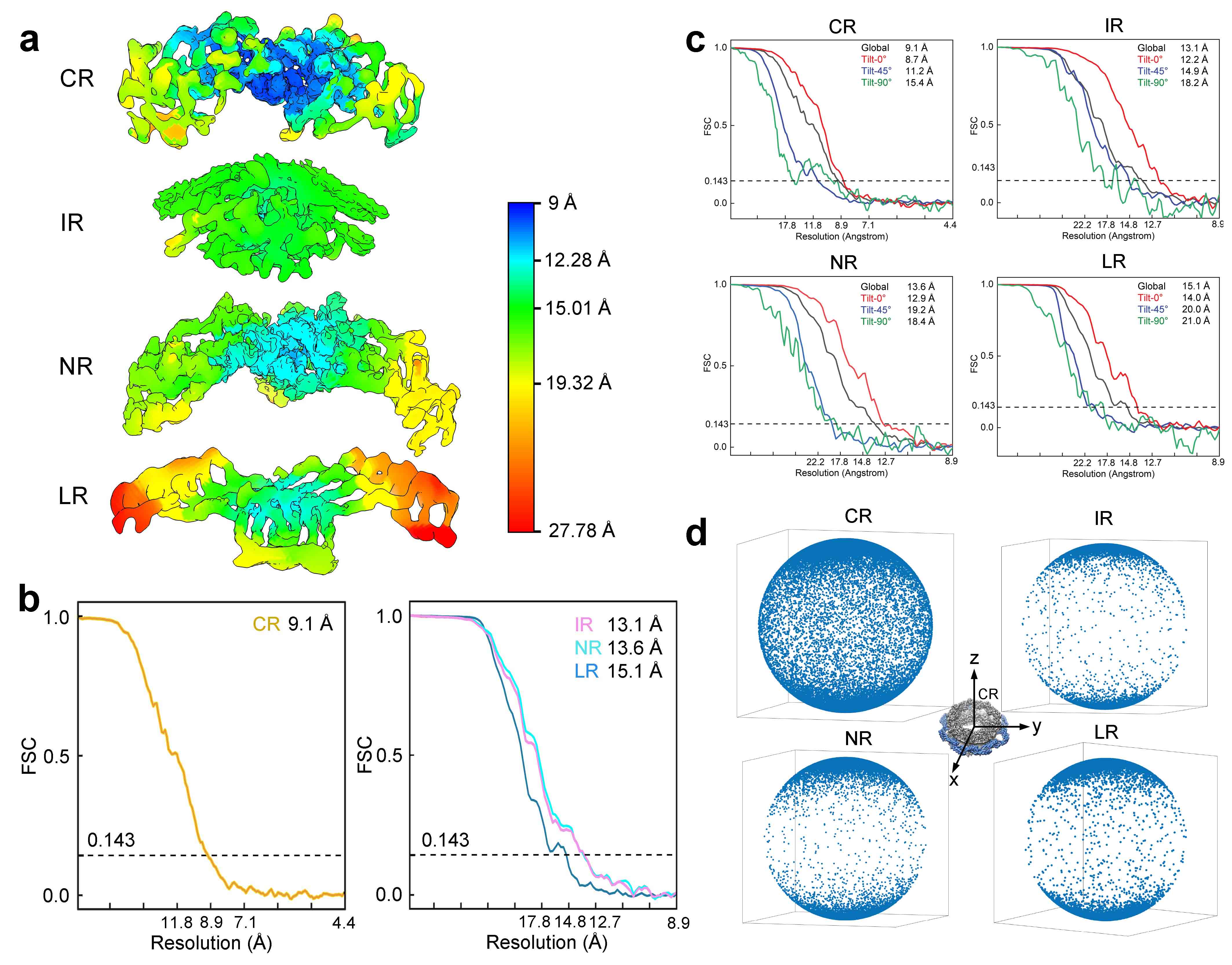

### Supplementary information, Fig. S4

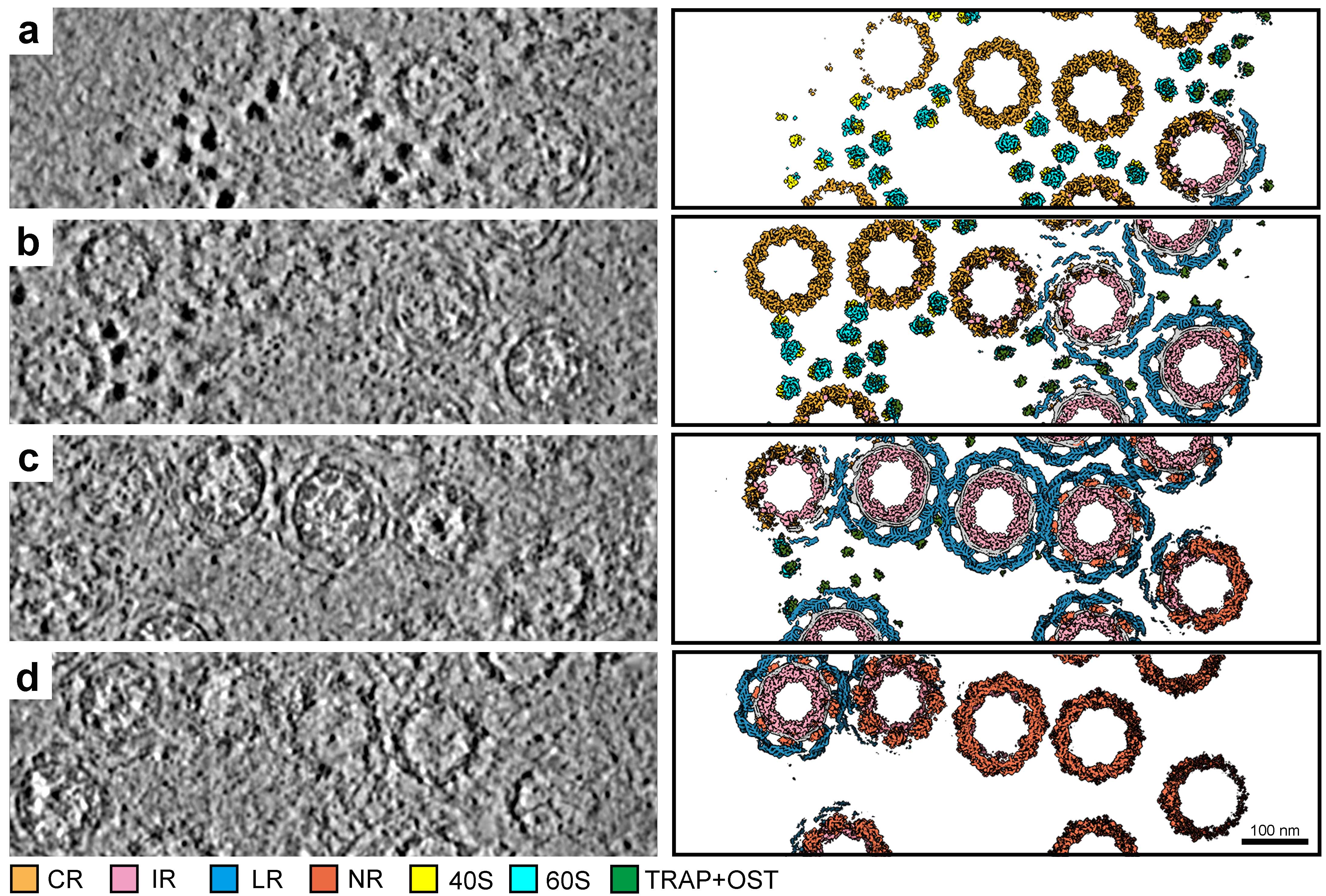

### Supplementary information, Fig. S5

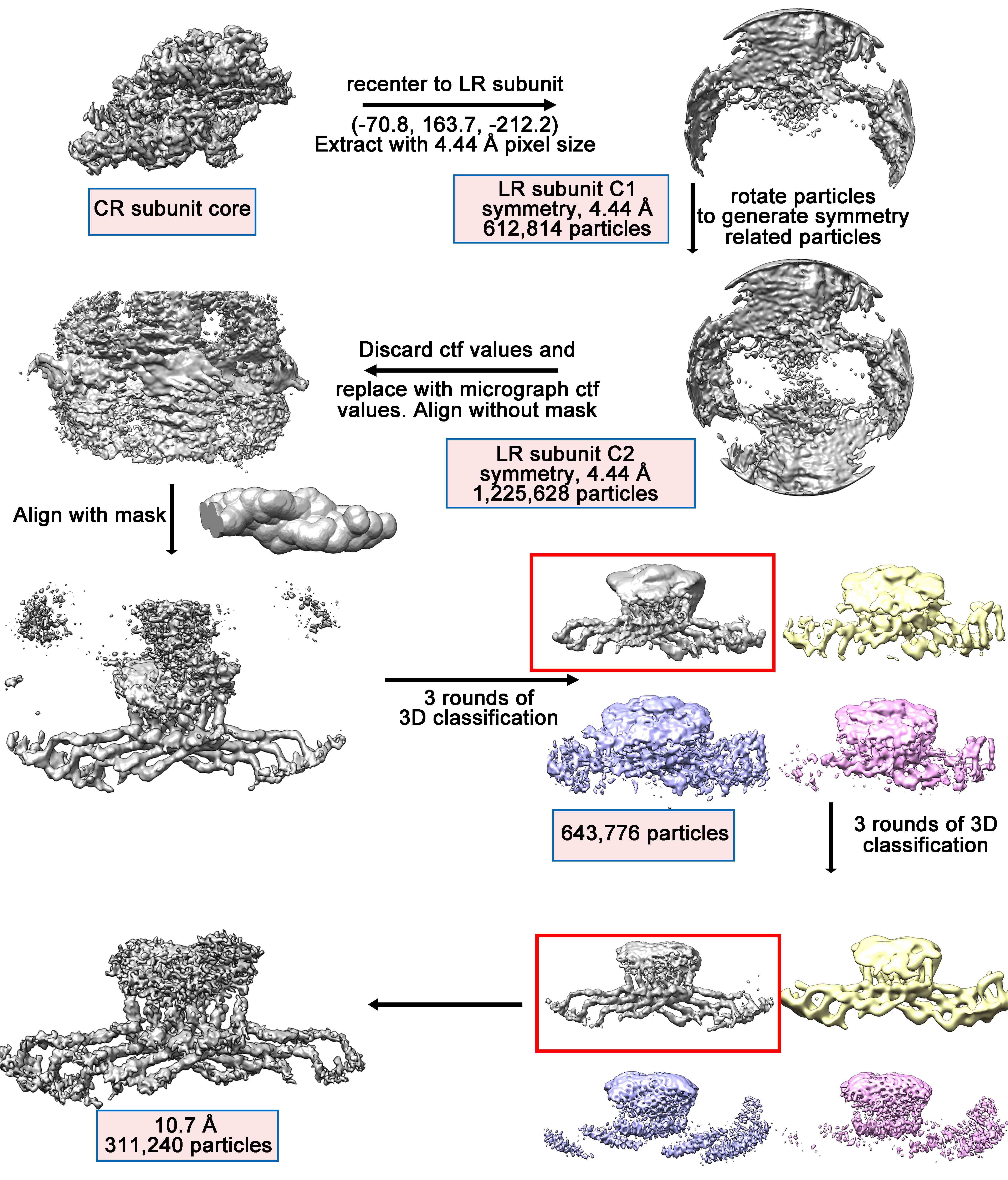

### Supplementary information, Fig. S6

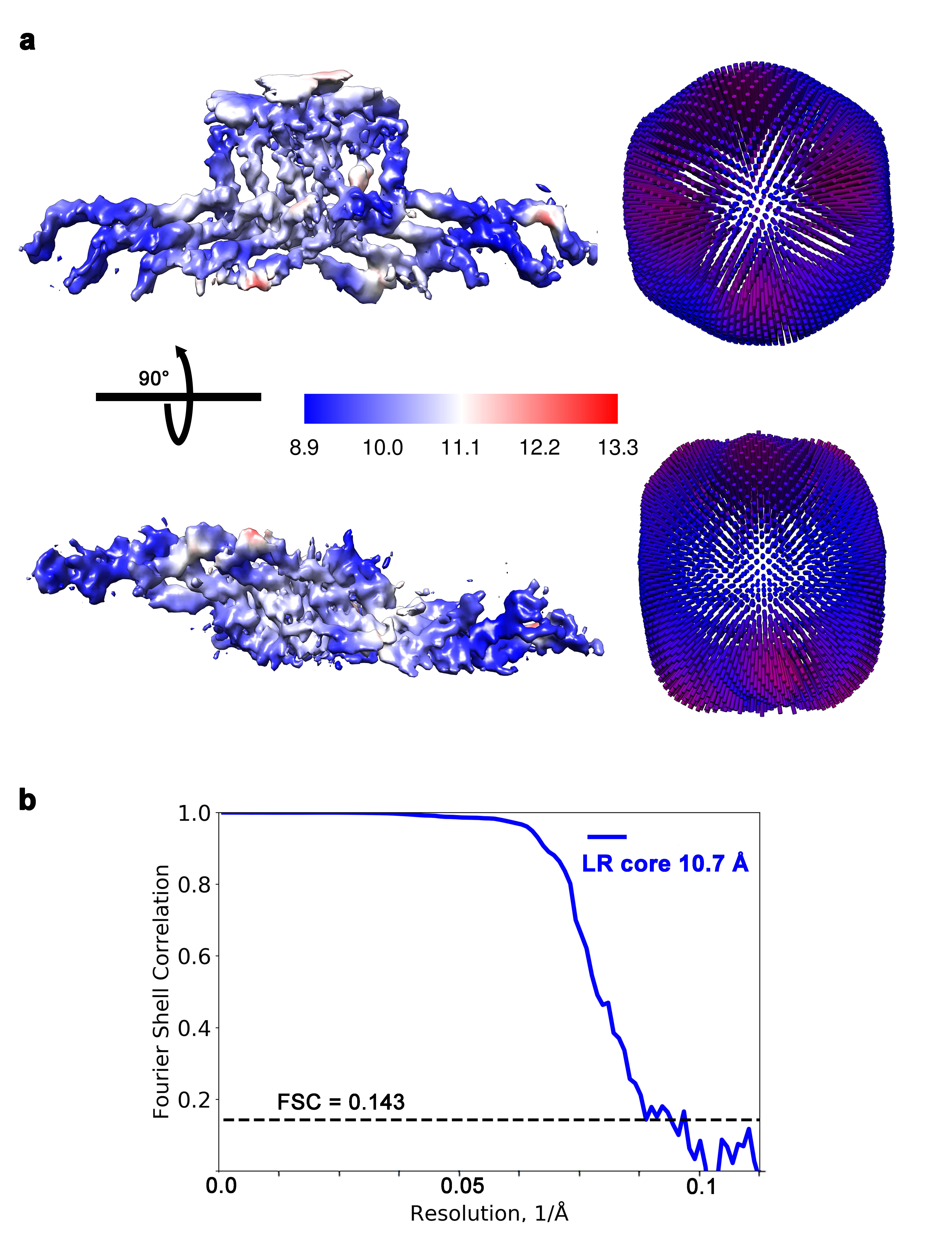

### Supplementary information, Fig. S7

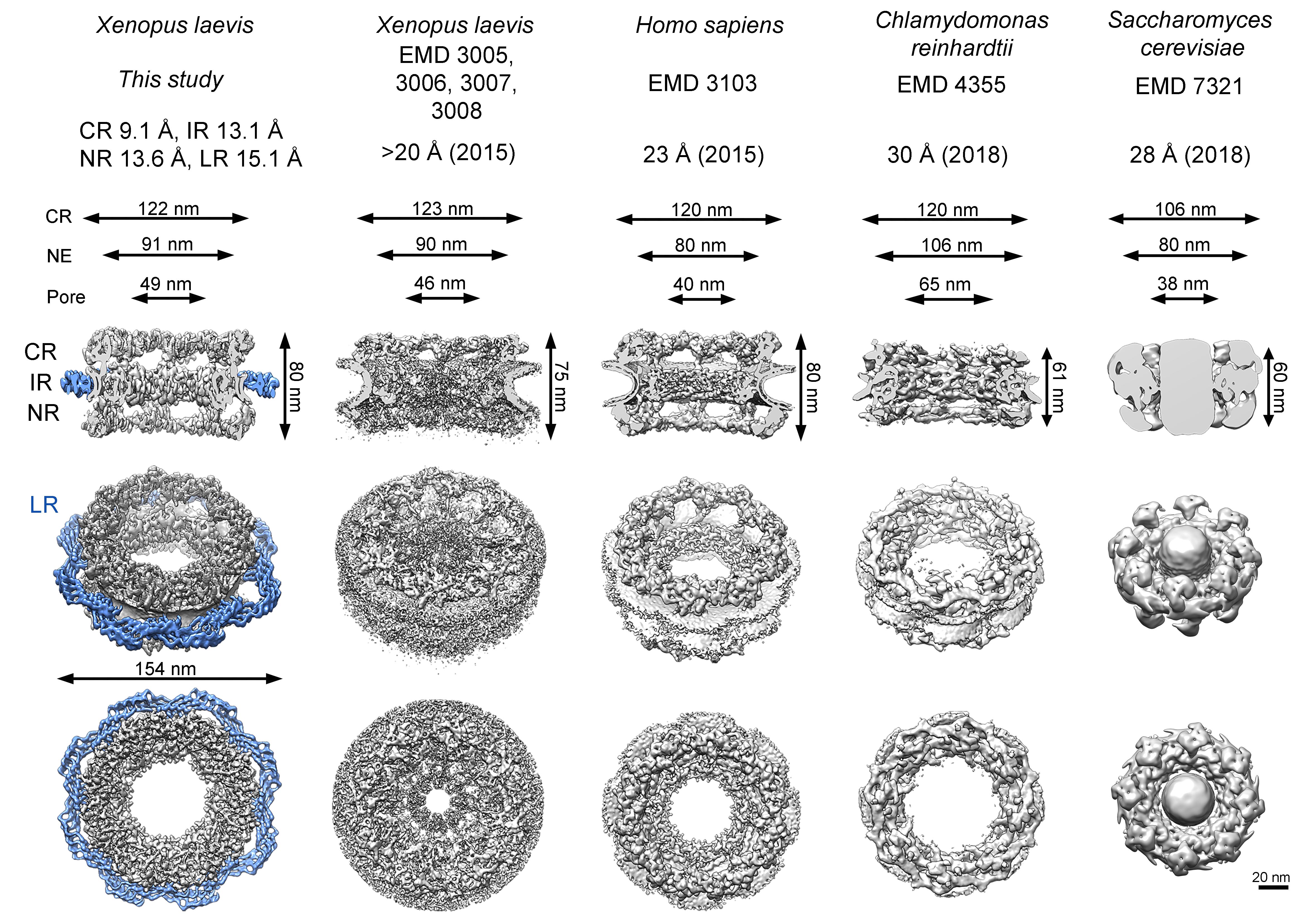

### Supplementary information, Fig. S8

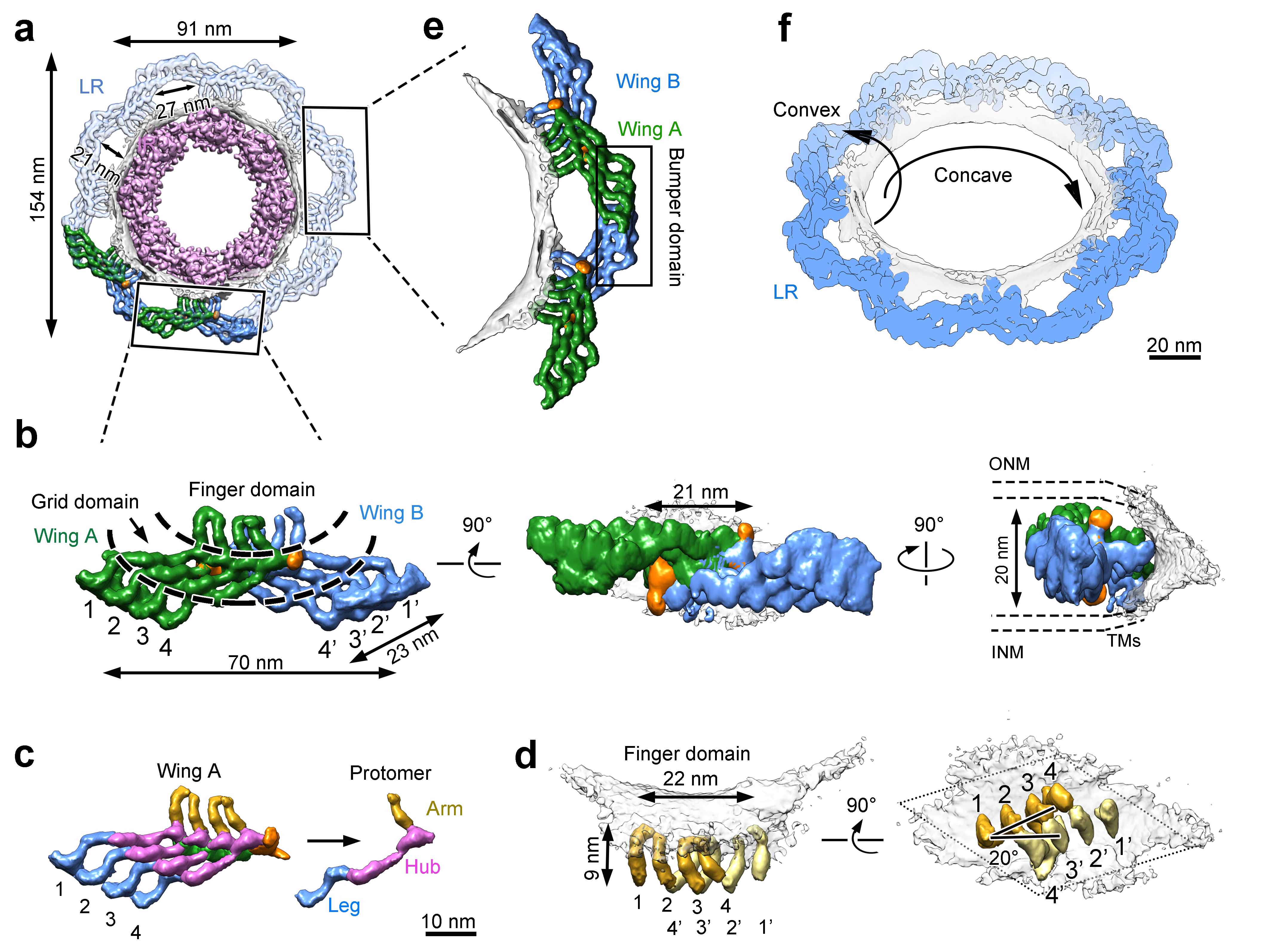

### Supplementary information, Fig. S9

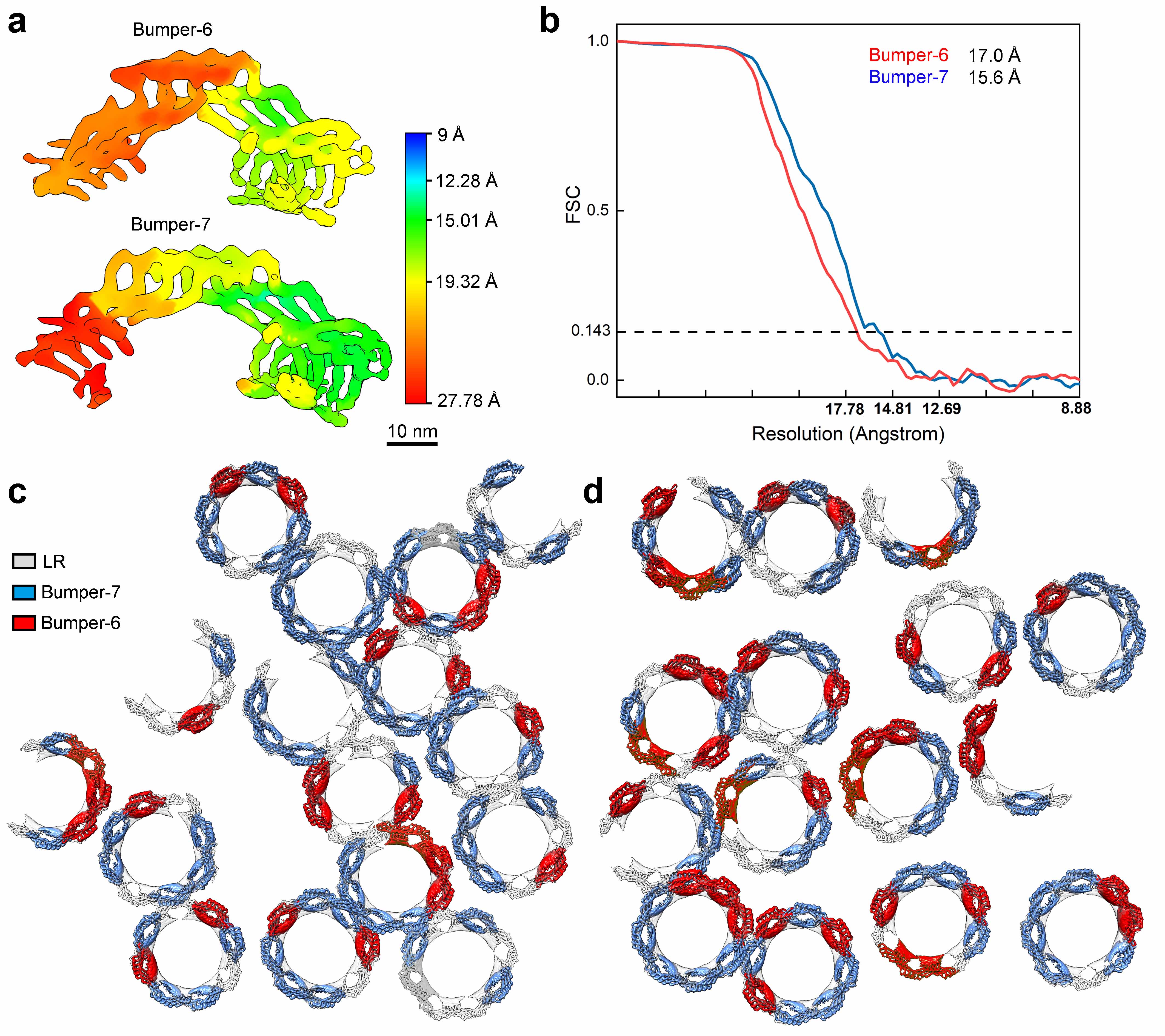

### Supplementary information, Fig. S10

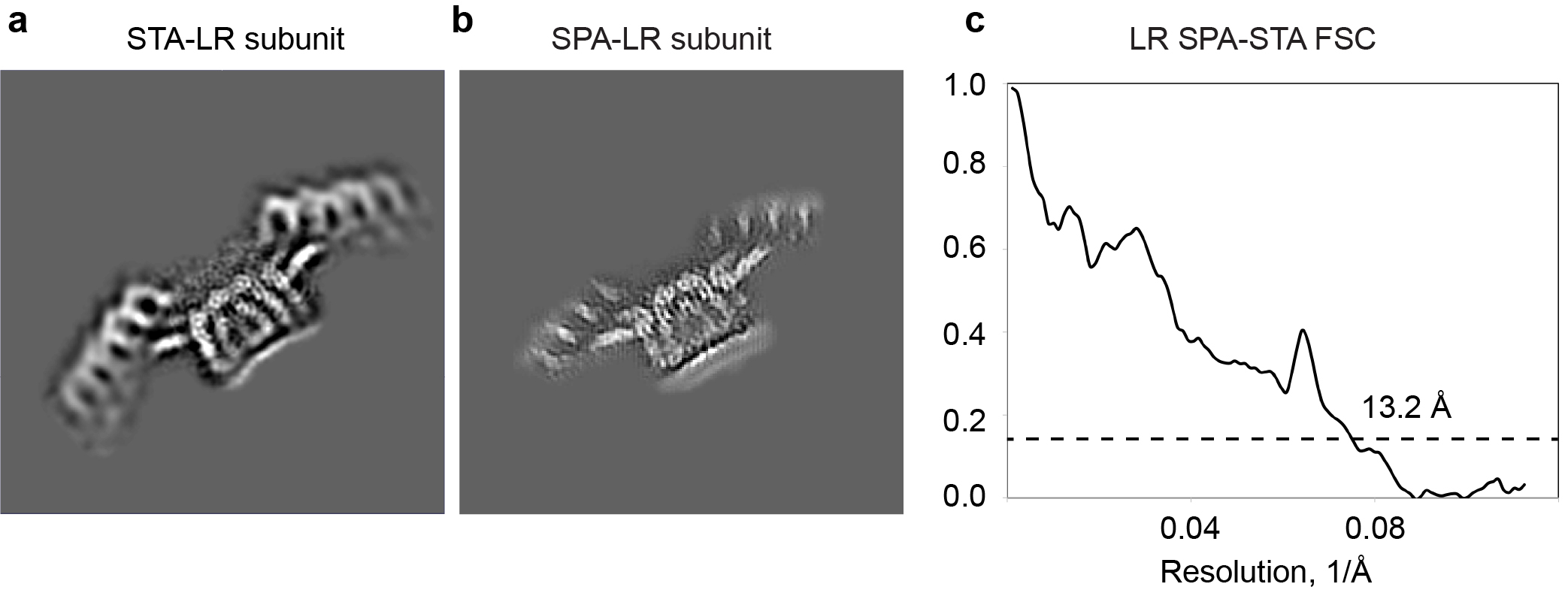

### Supplementary information, Fig. S11

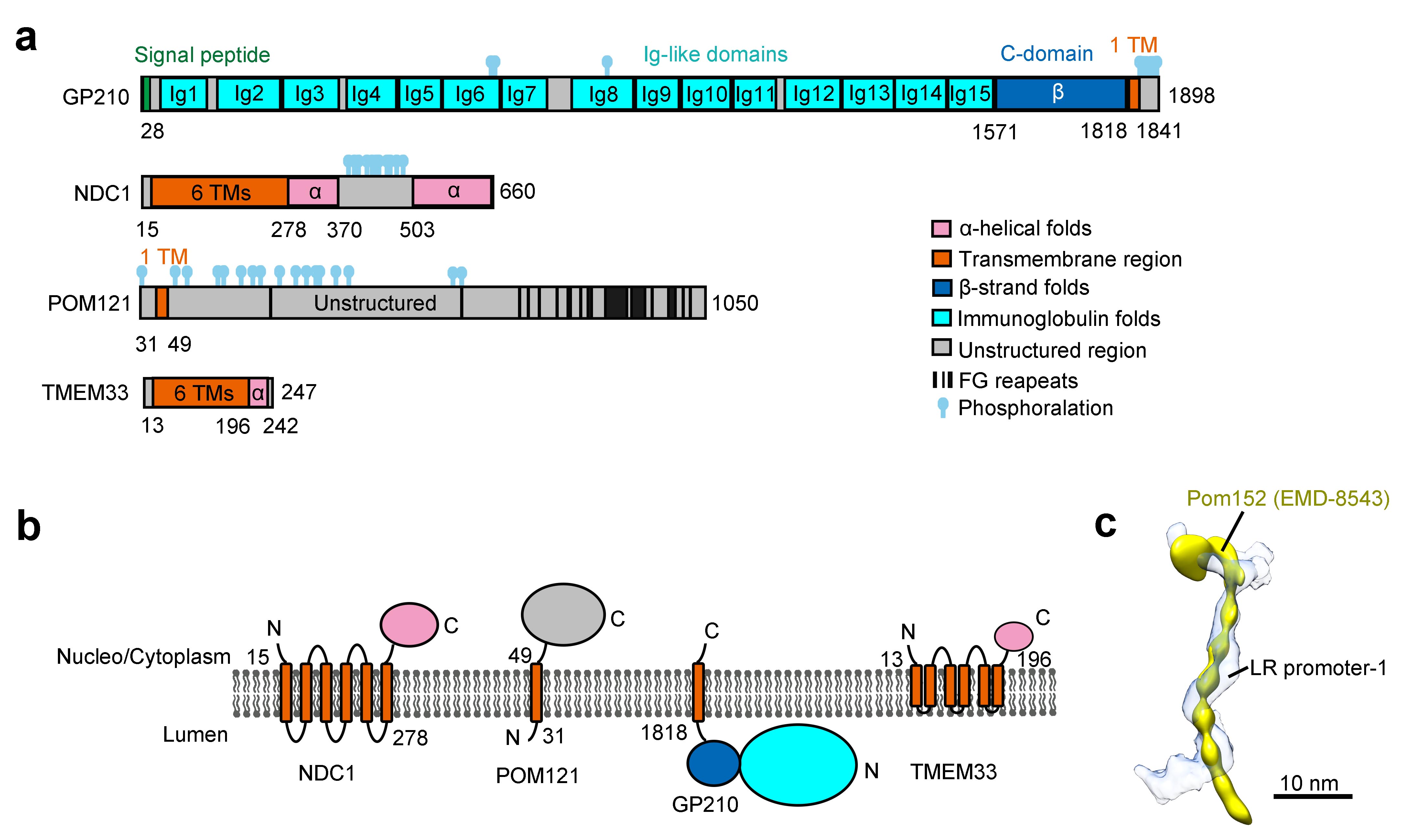

### Supplementary information, Table S1

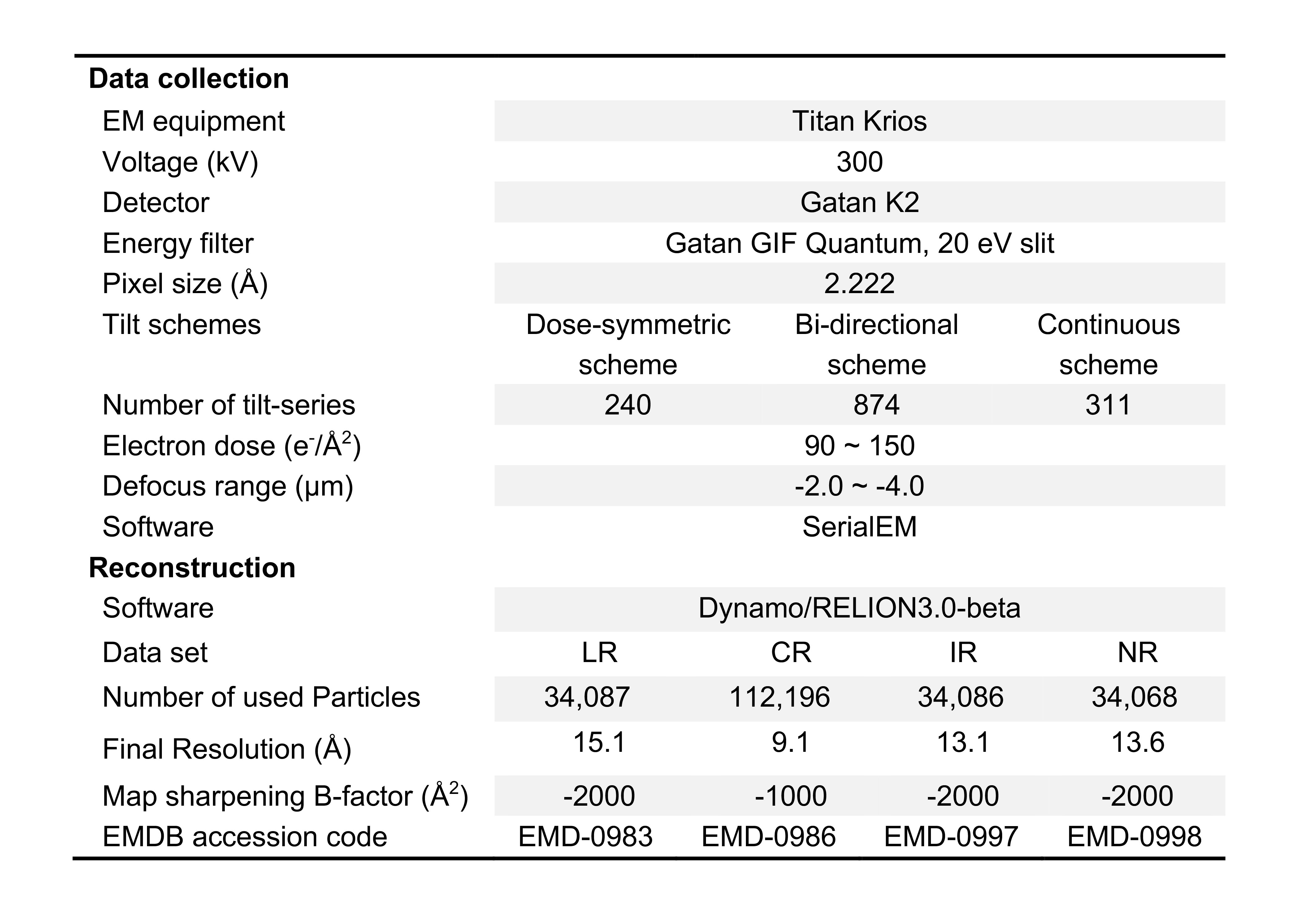

### Supplementary information, Table S2

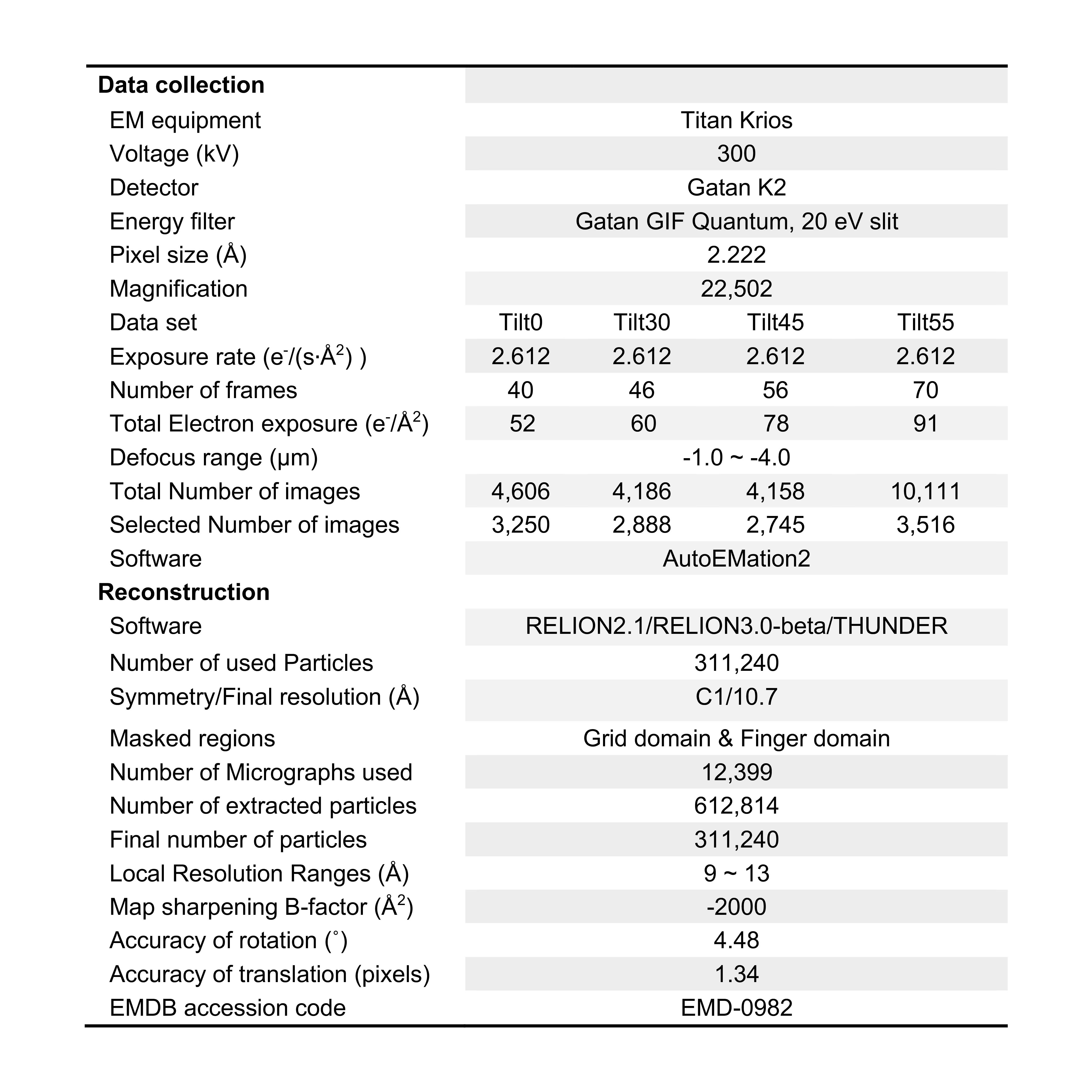
